# Nanoarchitecture of the presynaptic cell adhesion molecule PTPσ in relation to prospective vesicle fusion sites and postsynaptic NMDA receptors

**DOI:** 10.64898/2026.10.21.760724

**Authors:** Emily M. DeMarco Hibbs, Samira Mitias, Rebecca Aitken, Aaron D. Levy, Thomas A. Blanpied

**Affiliations:** Program in Neuroscience, University of Maryland School of Medicine, USA; Department of Pharmacology and Physiology, University of Maryland School of Medicine, USA; University of Maryland Medicine Institute for Neuroscience Discovery, USA

## Abstract

Protein organization at individual synapses tunes synaptic function. Presynaptic release-regulating proteins and postsynaptic receptors each concentrate into subsynaptic nanoclusters, and their precise alignment across the cleft into nanocolumns can maximize receptor activation following vesicle fusion. However, both the molecular organization defining presynaptic fusion sites and the mechanisms by which they are spatially coordinated with postsynaptic complexes remains unclear. Presynaptic transmembrane cell adhesion molecules may mediate both processes as they could simultaneously anchor cytosolic fusion machinery to the presynaptic membrane and bridge the cleft to coordinate PSD organization. The LAR-RPTP family member PTPσ is an important possibility, as its trans-synaptic interactors that bind PSD-95 are associated with NMDAR dysfunction, suggesting PTPσ may coordinate NMDAR positioning. However, PTPσ subsynaptic organization has not been determined. Here, we investigate the cellular expression and subsynaptic nano-organization of endogenous PTPσ in cultured rat hippocampal neurons. We find that PTPσ is enriched at most excitatory synapses, but unexpectedly, PTPσ nanoclusters do not colocalize with the vesicle priming protein Munc13-1. Instead, they adopt a stereotyped position adjacent to this machinery, without preferential association with sites in nanocolumns. PTPσ nanoclusters also lie offset from postsynaptic NMDA receptors, mirroring their relationship to vesicle fusion sites, yet the presence of PTPσ at a nanocolumn does not predict NMDAR abundance. Together, these findings suggest that PTPσ does not directly coordinate release site– NMDAR alignment but instead occupies defined nanoscale positions adjacent to vesicle fusion sites, potentially helping to shape or stabilize the vesicle priming landscape of the active zone.

**Significance statement:** Synaptic vesicles fuse at specific presynaptic locations, called release sites, whose precise position—and alignment with postsynaptic receptor density—critically regulates synaptic strength. NMDA receptors, which flux calcium to gate diverse forms of brain plasticity, are especially sensitive to glutamate release location. Here, we test whether the cell adhesion molecule PTPσ helps establish the relationship between release sites and NMDARs. We find that PTPσ occupies defined nanoscale positions adjacent to release sites, offering new insight into the molecular machinery organizing these sites and suggesting that PTPσ helps shape the probability distribution of vesicle fusion within the synapse.

## Introduction

Synaptic vesicle (SV) fusion with the plasma membrane is a key step in neurotransmission, and changes to the probability or abundance of SV fusion contribute to learning and disease (Alabi and Tsien, 2013; Liu et al., 2024). SV fusion in the presynaptic active zone (AZ) occurs only at a few discrete sites (Schikorski and Stevens, 1997; Imig et al., 2014) where the machinery necessary for vesicle fusion, including Munc13-1 and RIM1, accumulates into sub-100 nm protein densities termed nanoclusters (NCs) (Biederer et al., 2017). These concentrated proteins tether SVs at the membrane and couple them closely to voltage-gated Ca^2+^ channels (CaVs), enabling the formation of Ca^2+^ nanodomains that support rapid, Ca^2+^-triggered SV fusion (Nakamura et al., 2018; Rebola et al., 2019; Tan et al., 2022b). Thus, the precise molecular architecture of the presynaptic AZ dictates both the location and competence of SV fusion events.

SV fusion site positioning is a critical determinant of overall synaptic strength and signaling, because the distance between neurotransmitter release and postsynaptic receptors strongly influences receptor activation (Biederer et al., 2017). Similar to the AZ, postsynaptic neurotransmitter receptors (AMPA and NMDA receptors (AMPARs, NMDARs)) and the scaffolds such as PSD-95 that anchor them are organized into NCs (Fukata et al., 2024; Damenti et al., 2026). When presynaptic fusion machinery NCs are aligned across the synaptic cleft with these postsynaptic NCs in an arrangement called the nanocolumn (Tang et al., 2016), both NMDAR and AMPAR activation following AP-evoked vesicle fusion is maximized (Haas et al., 2018; Goncalves et al., 2020; Li et al., 2021; Ramsey et al., 2021; Anderson et al., 2025). Thus, the precise nanoscale architecture of synapses underlies efficient synaptic transmission and requires strong trans-synaptic coordination between the position of presynaptic SV fusion sites and postsynaptic receptors and scaffolds.

Despite the clear functional importance of AZ architecture, the mechanisms by which SV fusion complexes are anchored to the plasma membrane at specific sites within the AZ, and how these sites are spatially coordinated with postsynaptic complexes, remain unclear. Synapse-resident cell-adhesion molecules (CAMs) represent compelling candidates for mediating both processes (Dalva et al., 2000; Haas et al., 2018; Fukata et al., 2021; Ramsey et al., 2021; Dai et al., 2022; Liakath-Ali et al., 2022; Lloyd et al., 2023), as their membrane-bound nature could simultaneously anchor cytosolic fusion machinery to the presynaptic membrane and bridge the cleft to coordinate PSD organization. While several families of presynaptic CAMs form a multitude of cleft interactions with postsynaptic CAMs, only the LAR-RPTP family makes direct cytosolic interactions with the SV priming and fusion protein complex (Emperador-Melero et al., 2024); interactions made by the well-studied neurexins are a degree removed (Südhof, 2018). LAR-RPTPs directly bind the core AZ protein liprin-α, which recruits RIM1 and Munc13-1 to the AZ and chaperones SV priming (Wong et al., 2018; Wakita et al., 2020; Emperador-Melero et al., 2024; Marcó de la Cruz et al., 2024). Thus, this liprin-α-LAR-RPTP interaction has been proposed to be a molecular axis through which the SV priming complex is anchored to the plasma membrane and SV fusion sites are positioned (Emperador-Melero et al., 2024).

Of the three LAR-RPTP members, PTPσ is a noteworthy candidate to control excitatory synapse nanoarchitecture in the hippocampus. PTPσ expression persists at excitatory hippocampal synapses (Han et al., 2020) following its essential roles in synaptogenesis, synapse differentiation, and axon growth (Zhang et al., 1998; Thompson et al., 2003). Chronic PTPσ ablation significantly reduces SV fusion probability and increases presynaptic facilitation, suggesting disruptions to sites of evoked SV fusion (Han et al., 2020) and raising the possibility that PTPσ could coordinate the positioning of SV fusion machinery. Further, liprin-α3 causes RIM and Munc13-1 to form condensates in heterologous cells that only localize to the cellular membrane upon co-expression of PTPσ (Emperador-Melero et al., 2024), suggesting PTPσ can bridge release machinery to the membrane. However, the nanoscale distribution of PTPσ within the AZ, especially in relation to these other key proteins, is unknown.

PTPσ could also be a structural mediator of trans-synaptic alignment between SV fusion sites and receptors. First, its interactors NGL-3, TrkC and GPC/LRRTM4 all drive PSD-95 clustering, though direct involvement of PTPσ has not been tested (Wyszynski et al., 2002; Kwon et al., 2010; Takahashi et al., 2011; Ko et al., 2015; Li et al., 2015; Choi et al., 2016; Han et al., 2018; Roppongi et al., 2020; Khaled et al., 2024). Second, PTPσ may trans-synaptically regulate NMDAR function: most notably, presynaptic PTPσ ablation reduces synaptic NMDAR activation without changing receptor expression (Kim et al., 2020; Sclip and Südhof, 2020a), as would be predicted from receptor misalignment to SV fusion sites. These results suggest PTPσ may be preferentially positioned within the synapse to be near both NMDARs and SV fusion sites.

To investigate this possibility, we employed an array of confocal and super-resolution imaging approaches to explore the endogenous expression profile of PTPσ in established synapses. We found that PTPσ predominantly localized to the presynaptic AZ of most excitatory synapses and was subsynaptically organized into discrete NCs. Surprisingly, these NCs were not co-localized with SV fusion sites marked by Munc13-1 NCs but instead were systematically positioned adjacent to them but showed no preference for release sites associated with the nanocolumn. PTPσ NCs also showed a stereotyped relationship with NMDARs that was strikingly similar to their relationship with SV fusion sites, yet the presence of PTPσ was not predictive of NMDAR localization to sites of SV fusion or the nanocolumn. Together, our results suggest a model in vesicle fusion sites are assembled with PTPσ associated at the periphery where it could be ideally positioned to function in the vesicle priming and release complex of AZ proteins.

## Materials & Methods

### Animals

All animal procedures were approved by the Institutional Animal Care and Use Committee at the University of Maryland School of Medicine. Sprague-Dawley rats (Charles River) were housed individually before embryo retrieval at E18 for neuron culture. C57/BL6J mice bred in-house were weaned at postnatal day P21 and housed two to five per cage in single-sex groups. Adult mice were used for brain slice immunohistochemistry at P37. For both mice and rats, food and water were available ad libitum and lights were maintained on a 12 h light/dark cycle.

### DNA Constructs

pCAG GPHN.FingR-GFP-CCR5TC was a gift from Don Arnold (Gross et al., 2013) (Addgene plasmid #46295). psPAX2 (Addgene plasmid #12260) and pMD2.G (Addgene plasmid #12259) were gifts from Didier Trono. Lentiviral L-309 shPTPRS_UbC-IRES2-eGFP (shPTPRS) was a gift from Jaewon Ko (Han et al., 2018). LCV2-EGFP was described previously (Dharmasri et al., 2024a). pCMV_HA-tdTomato-hPTPRS was a gift from Pascal Kaeser. hEF1_mCherry-P2A-mPTPRD was generated by Golden Gate Assembly of three custom plasmids: a hEF1 promoter destination vector plus mCherry-P2A and PTPRD (synthesized from NM _011211.4; gift of Alexandros Poulopoulos) parts. CAG_mTagBFP2 was generated by Golden Gate Assembly from four custom plasmids: a CAG promoter destination vector plus Kozak-start codon, mTagBFP2, and stop codon parts.

### Cell culture

Dissociated hippocampal cultures were prepared from E18 Sprague-Dawley rats of both sexes as described previously (Dharmasri et al., 2024b) and plated on poly-L-lysine-coated coverslips (#1.5, 18 mm, Warner) at a density of 30,000 cells/coverslip for all experiments except when testing anti-PTPσ antibody specificity (Fig. S1A-D), where neurons were plated at 50,000 cells/coverslip then infected with 10-15 µl of shPTPRS lentivirus at DIV1. For all experiments, neurons were fixed for immunofluorescence staining at DIV19-21.

HEK293T cells (ATCC CRL-3216) were maintained in DMEM + 10% FBS and penicillin/streptomycin at 37°C and 5% CO^2^. For anti-PTPσ isoform specificity testing (Fig. S1E-F), HEK cells were plated on poly-L-lysine and fibronectin-coated 18 mm coverslips at a density of 120,000 cells/coverslip, transfected 24 h later with either 500 ng pCMV_HA-tdTomato-hPTPRS or hEF1_mCherry-P2A-mPTPRD + 150 ng CAG_mTagBFP2 with Lipofectamine 2000 for 4 hours, then maintained for 24 h in fresh media before fixation.

### Lentivirus

For lentivirus production, HEK293T cells were plated at 5×10^6^ cells/10 cm plate and transfected 12-24 h later with 6 µg of either shPTPRS or LCV2-EGFP + 4 µg psPAX2 + 2 µg pMD2.G using PEI for 4-6 hours. After 72 h, the media was harvested and debris removed by centrifugation at ∼170 x g for 5 min and 0.45 µm PES filtering. Single-use aliquots were frozen at -80°C for long-term storage without further concentration. Lentivirus routinely infected >90% of neurons at the volumes used.

### Antibody preincubation

We stained multiple targets in the same sample using antibodies from the same species by preincubating primary antibodies with secondary nanobodies as previously described (Pleiner et al., 2018; Sograte-Idrissi et al., 2020; Dharmasri et al., 2024b; Emperador-Melero et al., 2024; Metzbower et al., 2024; Anderson et al., 2025). We mixed each primary antibody with 3-fold molar excess of a secondary single-domain antibody (sdAb) labeled for either fluorescence (NanoTag Biotechnologies) or DNA-PAINT (Massive Photonics) and targeting the primary species for 20 minutes at room temperature (RT) in PBS + 100 mM glycine (PBS/Gly). Any sdAb “hopping” between primaries of the same species was prevented by addition of (4-fold) molar excess Multiplexing Blocker (NanoTag Biotechnologies) matched to the secondary sdAb target species for a further 20 minutes. Preincubated antibodies were then pooled together and diluted to their final concentrations in PBS/Gly.

### Brain slice immunohistochemistry

5-8 week old mice were anesthetized via intraperitoneal injection of ketamine (1.5 µg/mL) and xylazine (0.15 µg/mL) before being transcardially perfused with 2% paraformaldehyde (PFA). Brains were dissected and post-fixed for 30 minutes in the same fixative and cryoprotected in 30% sucrose/phosphate-buffered saline (PBS) before being flash-frozen in Optimal Cutting Temperature (OCT, Sakura) compound on dry ice and stored at –20°C. Brain sections were cut on a Leica CM1850 cryostat into 15 µm thick slices. Slices were mounted directly on (3-Aminopropyl)triethoxysilane (APTES)-coated coverslips and air-dried to promote adherence before being re-hydrated and stored in PBS at 4°C until permeabilization. Sections were permeabilized in 0.3% Triton X-100 in PBS for 20 minutes and then blocked with BlockAid Blocking Solution (Invitrogen) containing 0.2% Triton X-100 for 1h at room temperature. Sections were incubated with Atto488-labeled anti-PSD-95 sdAb (1:200; NanoTag Biotechnologies, N3702-At488-L, RRID: AB_3076105) and mouse anti-PTPσ primary antibody (1:400, MediMabs, MM-0020-P) preincubated with Alexa Fluor™ 647-labeled anti-Mouse sdAb (NanoTag Biotechnologies, N1202-AF647-S, RRID: AB_3075957) in BlockAid + 0.1% Triton X-100 overnight at 4°C. The following day, sections were rinsed 3x with PBS/glycine, and the coverslips were mounted on glass slides in Abberior Liquid Mount, sealed with nail polish, and stored at 4 °C.

### Immunostaining

All cultured neurons and HEK cells were fixed with 4% PFA in 4% sucrose and PBS for 8 min at room temperature (RT), washed in PBS/Gly 3 × 5 min at RT, permeabilized with 0.3% Triton X-100 for 20 min at RT, blocked with BlockAid + 0.2% Triton X-100 for 45 min at RT, labeled with primary antibodies preincubated with sdAbs in BlockAid + 0.1% Triton X-100 overnight at 4°C, washed, labeled with secondary reagents or additional sdAbs in BlockAid without Triton X-100 for 45 min at RT, washed, and postfixed for 15 minutes at RT.

For DNA-PAINT, DIV 19-21 neurons were labeled overnight with DNA-PAINT docking strand-conjugated sdAb-preincubated primary antibodies. DNA-PAINT sdAbs (Massive Photonics) carried one of four FAST DNA-PAINT strands (F1, F3, F4 or F6) and were preincubated with primaries as follows: mouse anti-PSD-95 (1:80; NeuroMab, 75-028, RRID: AB_2292909) + anti-Mouse sdAb F1; rabbit anti-Munc13-1 (1:250, Synaptic Systems 126103) + anti-Rabbit sdAb F4; mouse anti-PTPσ (1:400; MediMabs, MM-0020-P) + anti-Mouse IgG1 sdAb F6; rabbit anti-GluN1 (1:100; Sigma-Aldrich, G8913, RRID: AB_259978). Neurons were labeled the next day with Atto488-labeled anti-PSD-95 sdAb (1:250, NanoTag N3702-At488-L; RRID: AB_3076105) for visual identification of excitatory synapses.

For confocal, cells were stained overnight with mouse anti-PTPσ (1:400; MediMabs, MM-0020-P) preincubated with Alexa Fluor 647-labeled anti-Mouse sdAb (NanoTag Biotechnologies, N1202-AF647-S, RRID: AB_3075957) and the next day with Atto488-labeled anti-GFP sdAb (1:250; Nanotag Biotechnologies, N0304-At488-L, RRID: AB_2744629) and Alexa Flour 568-labeled anti-PSD-95 sdAb (1:250; NanoTag Biotechnologies, N3702-AF568-L; RRID: AB_3076103).

For STED, cells were stained overnight with Abberior STAR 635P-labeled anti-PSD-95 sdAb (1:500; NanoTag Biotechnologies, N3702-Ab635P-L; RRID: AB_3076102) and rabbit anti-Bassoon (1:50; Cell Signaling, 6897S) and either: mouse (IgG2B) anti-Bassoon (1:500; NeuroMab, 75-491), mouse (IgG2A) anti-Syntaxin 1A (1:500; Synaptic System, 110 111), mouse anti-PTPσ (1:400; MediMabs, MM-0020-P), Mouse (IgG1) anti-SAP102 (1:100; BioLegend, 832001) or Mouse (IgG1) anti-panShank (1:200; Antibodies Incorporated, 75-089, RRID: AB_10672418). The next day cells were stained again for anti-Mouse Alexa Flour 594 (1:500; Thermo Fisher Scientific, A-21203, RRID: AB_2535789) and anti-Rabbit Abberior STAR 460L (1:80; Abberior, ST460L-1002, RRID: AB_3720997). Coverslips were mounted on slides in Abberior Solid Antifade mountant.

All key primary antibodies were knockout/knockdown validated: anti-PTPσ in (Han et al., 2018) and in Supplementary Figure 1, anti-PSD-95 by the manufacturer, anti-Munc13-1 by multiple groups (Lai et al., 2017; Tan et al., 2022a), anti-GluN1 in (Dharmasri et al., 2024a).

### Confocal microscopy and analysis

Confocal images of brain slices were acquired on an Abberior Facility Line STED microscope equipped with an Olympus IX83 base and UPlanXApo 60x/1.42 NA oil immersion objective, 405, 485, 561, and 640 nm lasers for imaging, a pulsed 775 nm STED laser, and photon-counting APDs for emission detection. We identified the hippocampal brain region of tissue slices using PSD-95 nanobody fluorescence and anatomical landmarks and acquired 2-color confocal 2mm x 2mm tiled overviews of PSD-95 and PTPσ line-by-line with 5 µs pixel dwell time and 250 nm pixel size (pinhole 1.0 AU). Alexa Fluor 488 dye was excited by the 488 nm laser at 50% power with accumulation of 3 lines. Alexa Fluor 647 dye was excited by the 640 nm laser at 80% power with accumulation of 3 lines. Using the tiled overview, 12.5 x 12.5 μm regions were selected within each hippocampal layer to acquire 2-color confocal images of PSD-95 and PTPσ with a pixel size of 80nm.

Confocal images taken of primary cultures (Fig.1B) were acquired on a Nikon TI2 inverted microscope equipped with an Andor Dragonfly spinning disk confocal unit, a Plan Apo λD 60x/1.42 NA oil immersion objective. The excitation path (405/488/561/640) was supplied by an Andor ILE and was reflected to the sample through a 405/488/561/638 quadband polychroic (Chroma), and the emission light passed through the confocal unit and appropriate emission filters (ET525/50, ET600/50 (Chroma) or Em01-R442/647 (Semrock)) onto a Zyla 4.2+ sCMOS camera (Andor). Cultured neurons were imaged at 50% laser power and 200 ms exposure, and Z-stacks were acquired using a piezo fitted in the Nikon stage.

**Figure 1.**
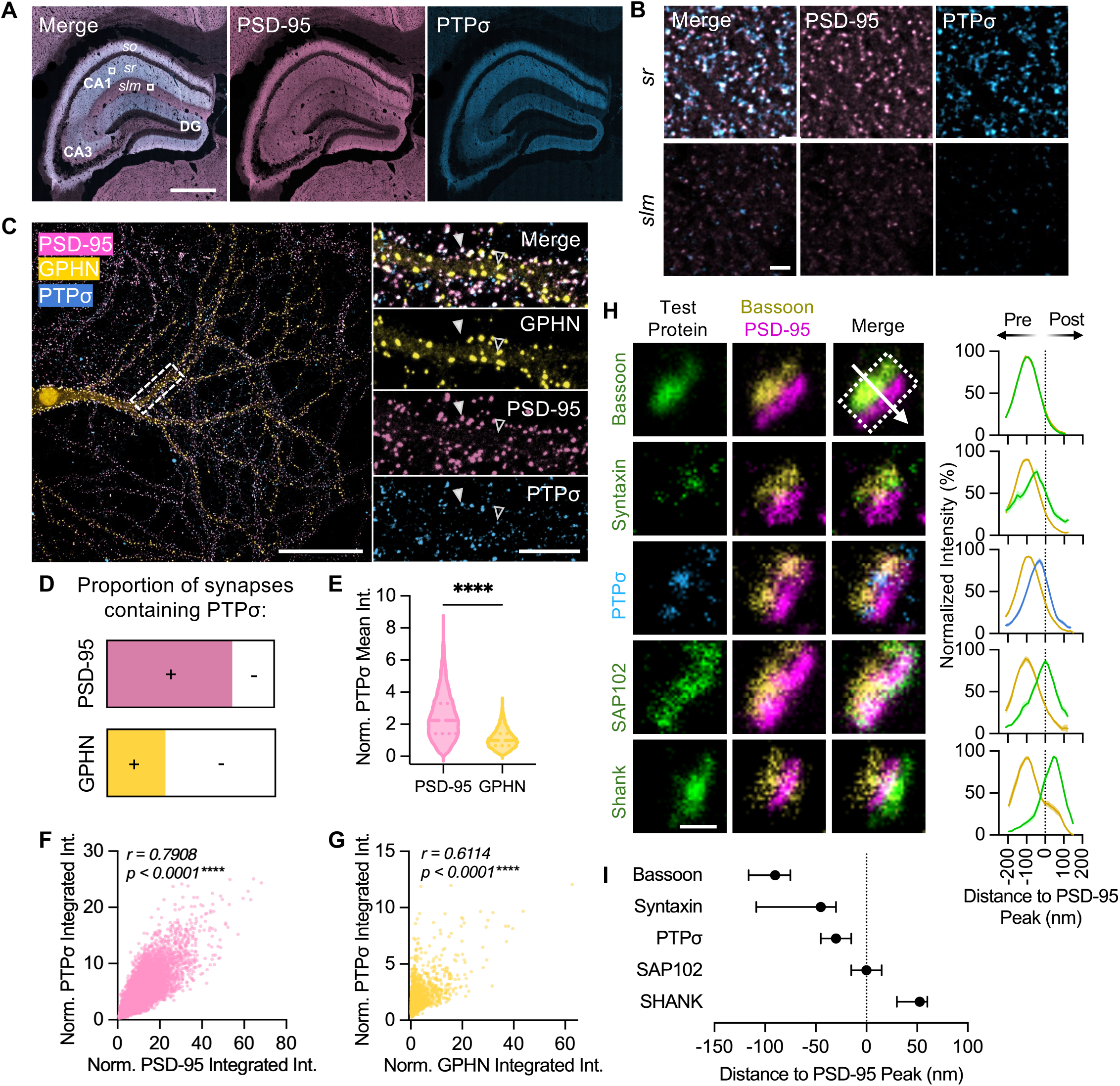
Endogenous PTPσ preferentially localizes to the presynaptic AZ of established excitatory synapses. **A**) Immunofluorescence images of PTPσ (blue) with PSD-95 (excitatory synapse marker; pink) in P37 mouse hippocampal slices. Hippocampal regions dentate gyrus (DG), CA3 and CA1 are indicated along with CA1 layers stratum oriens (*so),* stratum radiatum (*sr*) and stratum lacunosum-moleculare (*slm*). White boxes indicate locations of ROIs in (**B**). Scale bar 500 μm. **B)** Zoomed in view of boxed areas from **(A)** showing co-localization of PTPσ and PSD-95 in both *sr* and *slm,* but lower PTPσ levels in *slm* than *sr*. Scale bar 200 nm. **C**)Representative image of a GPHN.FingR (yellow)-expressing spiny neuron immunostained for PSD-95 (pink) and PTPσ (blue) in primary hippocampal neuron culture. White dashed box indicates inset region at right. (Insets) Solid arrowheads indicate PTPσ expression at an excitatory synapse and open arrowheads indicate PTPσ expression at an inhibitory synapse. Scale bar is 50 μm in overview and 10 µm in inset. **D**) Charts show the fraction of excitatory (PSD-95+, pink) or inhibitory (GPHN+, yellow) synapses with (colored) or without (white) PTPσ expression. **E**) PTPσ expression was significantly higher in excitatory synapses than inhibitory synapses. PTPσ mean intensity values were normalized to the median of the intensity values for a culture week (Mann-Whitney test P<0.0001). Violins show median (dashed line) and IQR (dotted lines). **F**) PTPσ expression showed a strong, positive correlation with PSD-95 expression (Pearson r = 0.7908, R^2^=0.63, P<0.001). **G**) PTPσ expression showed a weak, significant correlation with GPHN expression (Pearson r=0.6114, R^2^=0.37, P<0.001). Points in (**F-G**) are individual synapses. **H**) (Left) Representative STED images of side-view synapses for PTPσ (blue) or different test proteins (green; Bassoon, Syntaxin, SAP102 or Shank) at excitatory synapses identified by PSD-95 (magenta) and Bassoon (yellow). Scale bar 250 nm. Dotted box and arrow indicate example of size and direction of line scan used to generate normalized averaged intensity profiles (right) for each protein. Profiles were aligned to position the peak of PSD-95 at zero nm (dotted line). Lines show average intensity with standard error shading. (**I**) Quantification of the position of the peak intensity of each protein. Data are median +/- IQR. In **D-G**, n=19384 excitatory and 6745 inhibitory synapses across three independent cultures; in **H-I,** n = 36 Bassoon, 34 Syntaxin, 39 PTPσ, 28 SAP102 and 18 SHANK synapses. ***P<0.001; ****P<0.0001.

For excitatory/inhibitory synapse analysis (Fig. 1C-G), z-stacks were converted to maximum intensity projections using FIJI (Schindelin et al., 2012). Next, each image was cropped to isolate regions containing PSD-95 and GPHN.FingR signal on isolated dendrites to minimize background. Central regions were preferred to limit chromatic aberrations. Images were split into individual channels and background subtracted by the lowest 1% pixel value per channel. Putative synaptic puncta were identified independently for the channels containing PSD-95 and GPHN signals using the SynQuant plugin (Wang et al., 2020)(z-score 30 (PSD-95), 95 (GPHN); minfill 0.65; whratio 6; min size 10; max size 200). The shape and size of the resulting ROIs was then refined by dilating each ROI by 2 pixels to encompass the entire puncta before thresholding to 40% of the maximum intensity within the ROI, as we have previously described (Dharmasri et al., 2024b). For each punctum, area, mean intensity and integrated density were measured across all channels and exported as .csv files. The remainder of the analysis was performed using a custom MATLAB script. Likely mis-segmented ROIs were excluded from analysis based on the following measurements: 1) if an ROI was positive for both PSD-95 and GPHN (synapses should have only one or the other scaffold) – the cutoff for positive was set as the minimum mean intensity within SynQuant-segmented ROIs of that scaffold, and ROIs were excluded if they contained a greater mean intensity of the other scaffold than its cutoff; 2) if an ROI’s mean intensity in any channel was an outlier using the MATLAB ‘isoutlier’ function; 3) if an ROI’s area was less than 55 nm^2^ for both scaffolds or greater than 0.6 μm^2^ for PSD-95 ROIs only (GPHN ROIs tended to be more variable in size and substantially larger than PSD-95 overall, so were not subject to the same maximum size filter). Following filtering, the final dataset included 6745 inhibitory and 19384 excitatory synapse ROIs. To define PTPσ-positive and -negative synapses, the PTPσ channel image was rotated 180° and PTPσ background-subtracted mean intensity was re-measured within the same putative synaptic scaffold ROI. Since scaffold ROIs should be essentially randomly distributed relative to the rotated PTPσ image, the cutoff for an ROI to be PTPσ-positive was set at the highest non-outlier PTPσ mean intensity in the rotated image (where outliers are likely chance overlap of ROIs with PTPσ staining). Cutoffs were set independently for each culture to control for variations in staining or expression across cultures. Finally, intensity measurements for all proteins were normalized to the median intensity per culture.

### STED microscopy and analysis

STED images were acquired on the Abberior Facility Line STED microscope described above. We identified regions using PSD-95 and Bassoon fluorescence with 488 and 647 fluorescence emission then acquired 3-color STED images of PSD-95 (STAR 635P), Bassoon (STAR 460L) and each test protein antibody (Alexa Flour 594) line-by-line using 5 µs pixel dwell time and 15 nm pixel size (pinhole 0.91 AU). Abberior STAR635p dye was excited by the 640 nm laser at 25% power and depleted by the STED laser at 20% power, with accumulation of 5 lines. AlexaFlour594 dye was excited by the 561 nm laser at 70% power and depleted by the STED laser at 20% power, with accumulation of 5 lines. STAR460L dye was excited by the 488 nm laser at 75% power and depleted by the STED laser at 20% power, with accumulation of 5 lines. Corresponding confocal-resolution images were acquired prior to each STED image.

Data were analyzed using custom FIJI macros. For each STED image, side synapses were manually identified based as elongated PSD-95 puncta with an opposed but not overlapping Bassoon puncta, blind to the test protein signal. For each synapse, the integrated density of all three proteins (PSD-95, Bassoon and a test protein) was measured along a 35-pixel (525 nm) long line drawn perpendicular to the synaptic cleft with a width equal to that of the synapse. The remaining analyses were performed in GraphPad Prism. The intensity values for each protein were normalized to the peak intensity measurement within that synapse such that the peak intensity corresponds to 100% and the minimum intensity corresponds to 0%, and the distances for all three proteins were normalized to the position of PSD-95 peak intensity such that the PSD-95 peak corresponded to 0 nm.

### Single-molecule localization microscopy

DNA-PAINT images were acquired on a Nikon TI2 inverted microscope equipped with a 100x Apo TIRF oil-immersion objective (1.49 NA), 488, 561 and 640 nm excitation lasers routed through a TIRF illuminator, appropriate filters, a MicAO adaptive optics system (ImagineOptics), and a Zyla 4.2+ sCMOS camera (Andor). Hardware was controlled by Nikon Elements and MicAO software. Samples were imaged using HiLo illumination, and astigmatism for 3D imaging was applied using the MicAO and z-stacks of tetraSpeck beads (100 nm; Invitrogen) immobilized on separate coverslips were collected each imaging day for 3D calibrations. 90 nm gold nanoparticles were added to the sample before imaging at 1:10 dilution as fiducial markers.

Imaging regions were selected using PSD-95 nanobody Atto488 staining to identify synapse-dense dendrites of spiny, pyramidal-shaped neurons located within the TIRF illumination field. We imaged two regions per coverslip, across four culture weeks, and imaged four targets per region, exchanging solutions between each target (four exchange rounds total). The imaging buffer contained imager strands (Massive Photonics; F2-Cy3b, F3-Cy3b, F4-Cy3b or F6s-Cy3b) diluted in PBS + 500 mM NaCl pH 7.4 + 0.02% Tween-20 with oxygen scavengers (PCA/PCD/Trolox). Imager concentration was chosen independently for each target at each region, using the highest concentration that achieved the lowest background imager signal and minimized overlap between immobilized fluorescent events, concentrations ranging from approximately 100 to 500 pM. PTPσ was imaged first (F6s-Cy3b) and its localizations used to set the focal plane, then Munc13-1 (F2-Cy3b), PSD-95 (F4-Cy3b), and GluN1 (F3-Cy3b) were sequentially imaged. 25,000 images at 100 ms exposure were acquired for each exchange round, with output 561 nm laser power density at the objective of ∼170 W/cm^2^. After each round, imagers were exchanged by gently removing the imaging buffer and replacing it with 3 x 1 mL PBS + 500 mM NaCl.

### Single-molecule localization microscopy analysis

DNA-PAINT videos were localized with Picasso (Schnitzbauer et al., 2017) using a minimum net gradient 9,000 to detect molecules. The resulting localization files were independently drift-corrected with Picasso’s undrift by AIM function (Ma et al., 2024) using default parameters. Exchange rounds were aligned in Picasso first by RCC (Wang et al., 2014) followed by additional alignment using in focus gold fiducials and Picasso’s “undrift by picked” function.

Putative “en face” synaptic clusters of Munc13-1 and PSD-95 were selected in Picasso using the “pick” function, then exported to MATLAB for further refinement. Synapses were retained for analysis only if there were truly en face, i.e. imaged along the optical imaging axis (as determined by visual identification of synaptic cleft orientation during 3D rotation), had comparable sampling density for Munc13-1, PSD-95 and GluN1, and lacked obvious visual aberrations in any channel. Localization clusters that clearly came from non-synaptic regions within the original, circular picked region (which was larger than the actual synapse) were manually excluded. Example overview images were generated using the Fiji ThunderSTORM (Ovesný et al., 2014) plugin’s Averaged Shifted Histogram method with 8x magnification (20 nm pixels).

Nanoclusters were detected independently for each protein and synapse using Picasso’s HDBSCAN function (Campello et al., 2013) using 75 minimum cluster size, 10 (for PSD-95) or 5 (for Munc13-1, GluN1 and PTPσ) minimum samples and 130 inter-cluster max distance points. Only synapses containing at least one PTPσ NC were included in further analyses (N = 95). NC area outliers were identified and removed with GraphPad Prism’s ROUT method, using conservative parameters (Q=0.1%). The center of each NC was generated by Picasso’s HDBSCAN function. The diameter of each NC was calculated derived from a 2D alpha shape that completely enclosed all NC localizations, when approximating its shape as a circle.

Calculations of NC positioning relative to synapse center and periphery were performed as described in Anderson et al (2025). All protein localizations for a given synapse were combined and fit to an ellipse to derive a concentric ellipse with the center of a NC on its perimeter. The ratio was calculated for the area of the two ellipses, which yielded a two-dimensional measure of how close a given NC is located to the synapse center or edge. A linear representation of this was derived by taking the square root, where 0 is a NC at the center of the synapse, 1 is a NC on the edge of the synapse, and 0.5 is a NC positioned halfway between center and edge.

NC pairing was performed for each PTPσ and Munc13-1 NC by identifying, for each protein the nearest NC of the other protein and assessing whether those two NCs are mutually closest to each other, as described in Anderson et al (2025).

Auto- and cross-enrichments were analyzed in 2D using custom MATLAB functions similar to previously described (Dharmasri et al., 2024b; Anderson et al., 2025). In general, enrichments were measured by calculating the normalized density of a protein’s localizations as a function of distance from either the same (auto) or a different (cross) protein’s NC center. Enrichments were normalized to a random distribution of the measured protein’s localizations and smoothed by replacing high outliers (identified using Prism’s ROUT with Q = 0.1%) with the highest non-outlier value at each distance bin. Because of our high-resolution DNA-PAINT imaging, many apparent small clusters of localizations were likely to result from repeated localizations at a single docking strand, so randomizing all points would not yield a comparable distribution. Therefore, randomized distributions were created by first identifying putative single binding sites using Picasso’s SMLM clusterer function (Schlichthaerle et al., 2021; Reinhardt et al., 2023) which predicts subsets of localization point clouds that may originate from a single DNA strand. Then, the centroid position of each putative binding site was randomized within the bounds of the alpha shape of that protein, such that all localizations belonging to that site were moved together. Finally, we added a number of randomly placed localizations within the same alpha shape equivalent to the number of localizations not designated as part of a cluster. For PTPσ, we used the combined PTPσ and Munc13-1 alpha shape as PTPσ localizations were insufficiently abundant, and we expected PTPσ to fall within the AZ. To avoid any instability stemming from single randomizations of sparsely distributed proteins, each NC’s enrichment is reported as the average following 20 separate randomizations. To test whether a distribution was random (or not stereotyped), we also calculated selected cross-enrichments relative to random positions within the synapse, rather than real NC centers.

The enrichment index was calculated for GluN1 by averaging its enrichment within 60 nm of each Munc13-1’s NC center, as previously described (Anderson et al). We identified nanocolumnar Munc13-1 NCs essentially as described (Anderson et al) by comparing the enrichment index of each Munc13-1 NC with PSD-95 to that following 50 randomizations of PSD-95’s distribution. Munc13-1 NCs were considered enriched (in nanocolumn) or de-enriched (out of nanocolumn) when the real enrichment index of PSD-95 was greater or less than the average ±1.96 standard deviations of the enrichments to the 50 random distributions, respectively.

### Experimental Design and Statistical Analysis

Group measurements are expressed as median±IQR unless otherwise indicated. NC numbers were compared using Wilcoxon matched-pairs signed rank test. NC diameters and enrichment indices were compared using Mann-Whitney tests. Correlation comparisons were estimated using Pearson correlation coefficients.

## Results

### PTPσ is a presynaptic AZ protein primarily expressed at excitatory synapses

Before evaluating PTPσ nanostructural organization, we sought to confirm the presence of endogenous PTPσ in established hippocampal synapses (Han et al., 2018; Sclip and Südhof, 2020b), after its known roles in synaptogenesis and development (Takahashi et al., 2011; Takahashi and Craig, 2013; Um and Ko, 2013; Li et al., 2015; Khaled et al., 2024). We first re-validated the specificity and selectivity of a commercial antibody targeting PTPσ. Anti-PTPσ signal was lost after PTPσ knockdown in cultured primary hippocampal neurons (Fig S1A-D), replicating previous results (Han et al., 2018). We also demonstrated this antibody lacks reactivity against PTP∂, the other hippocampus-expressed LAR-RPTP family member (Han et al., 2018)(Fig. S1E-F). We immunostained P37 mouse hippocampal brain slices using this antibody and found that PTPσ expression was abundant across CA1 neuropil layers and concentrated in excitatory synapses, as indicated by co-localization with PSD-95 (Fig. 1A-B). PTPσ similarly was expressed at PSD-95-marked excitatory synapses in DIV21 cultured rat hippocampal neurons (Fig. 1C), a time point after synapses are established. These results indicate endogenous PTPσ remains present in established synapses well after synaptogenesis, both in vivo and in cultured neurons.

Interestingly, we did observe some PTPσ puncta co-localized with inhibitory synapses in culture that were marked by Gephyrin intrabody expression (GPHN, inhibitory post-synaptic scaffold) (Fig. 1C, open arrow). To directly assess their respective PTPσ abundance, we compared PTPσ content in excitatory synapses marked by PSD-95 immunostaining to inhibitory synapses marked by GPHN intrabody expression (Fig. 1C). PTPσ was present at ∼75% of PSD-95 puncta, consistent with previous results (Han et al., 2018), but was also present at ∼35% of GPHN puncta (Fig. 1D), indicating that while PTPσ is primarily expressed at excitatory synapses, it can also be expressed at some inhibitory synapses, at least in neuron culture. When expressed, PTPσ expression levels were also different between synapse types: PTPσ content was higher within excitatory synapses than inhibitory (Fig. 1E), and PTPσ content was strongly correlated with PSD-95 content (Fig. 1F, r=0.79, R^2^=0.63, p-value<0.001**) but less so (though still significantly) with GPHN (Fig. 1G, r=0.51, R^2^=0.37, p-value<0.001**), suggesting PTPσ could play unique roles at each synapse type. Thus, our data support previous literature that PTPσ is primarily expressed at excitatory synapses in established rat hippocampal culture (Han et al., 2018), enabling a deeper investigation of its distribution within these synapses.

PTPσ is known to enrich at the presynaptic AZ (Emperador-Melero et al., 2024) and in growth cones (Zhang et al., 1998), but biochemical data from both central and non-central synapse types suggest it could also be expressed at the post-synapse (Wyszynski et al., 2002; Dunah et al., 2005; Horn et al., 2012; Avilés et al., 2024). We therefore tested endogenous PTPσ localization to the pre- vs postsynaptic compartment of excitatory synapses in our cultured hippocampal neuron system using a variant of a well-established super-resolution microscopy assay of protein distribution along the trans-synaptic axis (Dani et al., 2010; Wong et al., 2018; Emperador-Melero et al., 2024). We immunostained DIV21 cultured hippocampal neurons for PSD-95 and the AZ marker Bassoon as post- and presynaptic landmarks, and co-stained for either PTPσ or one of a panel of control proteins with different known synaptic localizations: a different epitope on Bassoon, the presynaptic membrane-localized SNARE syntaxin-1a, the postsynaptic membrane-proximal scaffold SAP102, or the membrane-distal postsynaptic scaffold Shank. We then imaged side-oriented synapses with 3-color STED microscopy (Fig. 1H) and determined the pre- vs post-synaptic localization of each protein by measuring the peak position of the intensity distribution relative to the reference PSD-95 and Bassoon. As expected, the intensity peaks of the control proteins were all positioned near their reference pre- and postsynaptic markers, with Shank positioned deeper in the PSD than both SAP102 and PSD-95, which were positioned similarly (Fig. 1I). PTPσ’s position was consistent with that of a presynaptic transmembrane protein: it was overall more strongly co-localized with the presynaptic versus the postsynaptic compartment, the position of its intensity peak ∼30 nm from the PSD-95 peak was comparable to the position of known presynaptic membrane-localized syntaxin-1A, and this distance exceeded the expected distance to the postsynaptic membrane (Jeyifous et al., 2016). These data together indicate endogenous PTPσ is preferentially localized to the presynaptic AZ of established excitatory hippocampal synapses, well after its known roles in development.

### PTPσ organizes into few discrete nanoclusters at excitatory synapses

Having demonstrated PTPσ expression in established synapses, we next set out to measure PTPσ nanostructure and its sub-synaptic spatial relationships with pre and postsynaptic proteins implicated in AZ organization and NMDAR signaling. We quadruple-immunostained DIV 21 rat primary hippocampal neuron cultures for DNA-PAINT of PTPσ along with Munc13-1 as a marker of release sites (Böhme et al., 2016; Reddy-Alla et al., 2017; Sakamoto et al., 2018; Midorikawa et al., 2024), the obligate NMDAR subunit GluN1, and PSD-95, which we use along with Munc13-1 to define the nanocolumn (Dharmasri et al., 2023; Anderson et al., 2025). We performed 3D DNA-Exchange PAINT microscopy to map the nanoscale relationships of all four proteins at single synapses (Fig. 2). Individual synapses were identified based on coincident PSD-95 and Munc13-1 localization clusters, and we restricted analysis to synapses with abundant PTPσ (∼73% of the imaged population, consistent with the confocal analysis (Fig 1D), see Methods for details). We further limited our analysis to synapses with the cleft oriented perpendicular to the optical axis (en face), enabling the highest resolution for measuring distances between proteins across the face of the synapse. We then progressively analyzed different protein relationships within this set of synapses.

**Figure 2.**
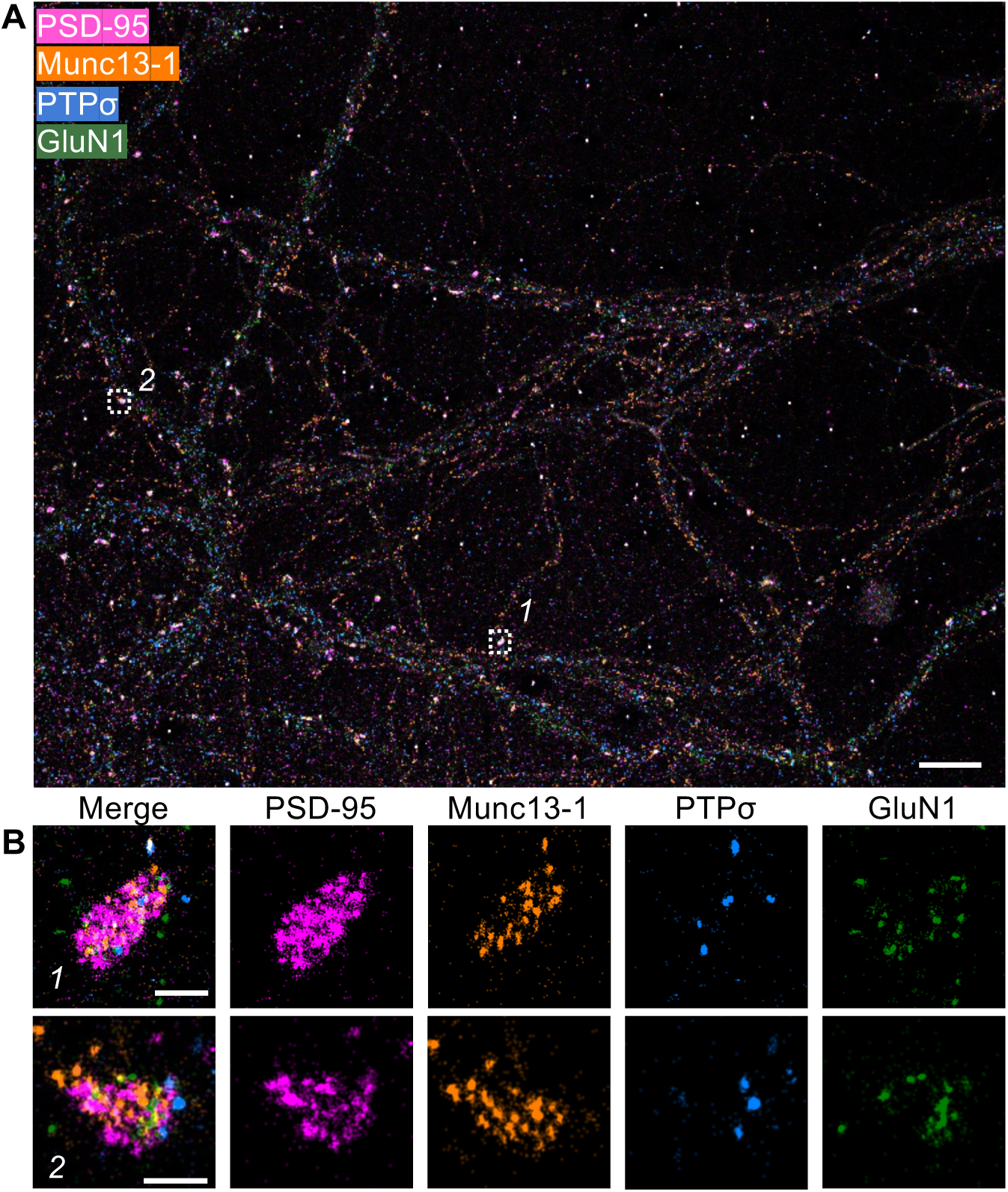
Mapping endogenous PTPσ organization using 3D DNA-Exchange PAINT. Raw DNA-PAINT rendering of primary rat hippocampal neuron culture at DIV21 immunostained for PSD-95 (magenta), Munc13-1 (orange), PTPσ (blue) and GluN1 (green). White boxes indicate locations of ROIs in (**B**). Scale bar 5 µm. **B**) Zoomed in view of boxed regions in (**A**) showing representative en face synapses. Localizations have been limited to those within +/-120 nm of the focal plane to aid visualization. Scale bar 250 nm.

We began by analyzing the subsynaptic organization of endogenous PTPσ itself. PTPσ was found almost exclusively in discrete, concentrated areas of high localization density (nanoclusters, NCs) that were readily discernible by eye in reconstructed images (Fig. 3A). We noted that similar NCs were also visually apparent in high-resolution STED images of en face synapses (Fig. 3B) taken from the dataset in Fig. 1H. We identified NCs in the DNA-PAINT data using HDBSCAN (Hierarchical density-based spatial clustering of applications with noise (McInnes et al., 2017), see Methods). Given the highly punctate nature of PTPσ signal, the number and size of PTPσ NCs were relatively insensitive to HDBSCAN parameter choices. We selected parameter values that minimized the detection of spurious localizations and minimized joining of intensity peaks separated by visible troughs. Parameters were chosen for other proteins based on prior work, which yielded a similar number of NCs per synapse as in those publications (Kellermayer et al., 2018; Sakamoto et al., 2018). We detected an average of 3.2 ± 0.2 PTPσ NCs per synapse (Fig. 3C) and the number of PTPσ NCs per synapse was positively correlated with AZ size (using Munc13-1 localizations to define the AZ borders) (*r* = 0.53, R^2^=0.28, *p* < 0.001**; Fig. 3D), though their number remained strikingly variable among AZs of similar sizes. PTPσ NCs had an average diameter of 70 ± 1.7 nm when approximating a circular NC (Fig. 3E). We confirmed this size through an autoenrichment (AE) analysis (Tang et al., 2016; Chen et al., 2020; Dharmasri et al., 2023), which measures localization density as a function of distance from the center of a given NC, normalized to the density predicted if the protein were randomly distributed within the bounds of the synapse. Consistent with the radius of HDBSCAN-detected NCs (∼70 nm diameter = 35 nm radius), PTPσ AE showed an elevated density at the center of a NC and maintained enrichment greater than random for 35 to 45 nm (Fig. 3F) and was not observed around random positions within the synapse. We also assessed the relative positioning of PTPσ NCs from the center to the edge of the synapse, normalizing the distance such that zero indicates a position in the center of the synapse and 1 at the edge (Anderson et al., 2025). PTPσ NCs were preferentially localized closer to the edge of the synapse compared to randomized positions, with only 40% of PTPσ NCs localized centrally and the remaining 60% localizing near the periphery. Additionally, 2% of the NCs were positioned outside of the synapse borders (Fig. 3G), suggesting a small extra-synaptic population may exist. Taken together, PTPσ forms small, discrete NCs that scale in number with AZ size and exhibit distinct positioning within that space.

**Figure 3.**
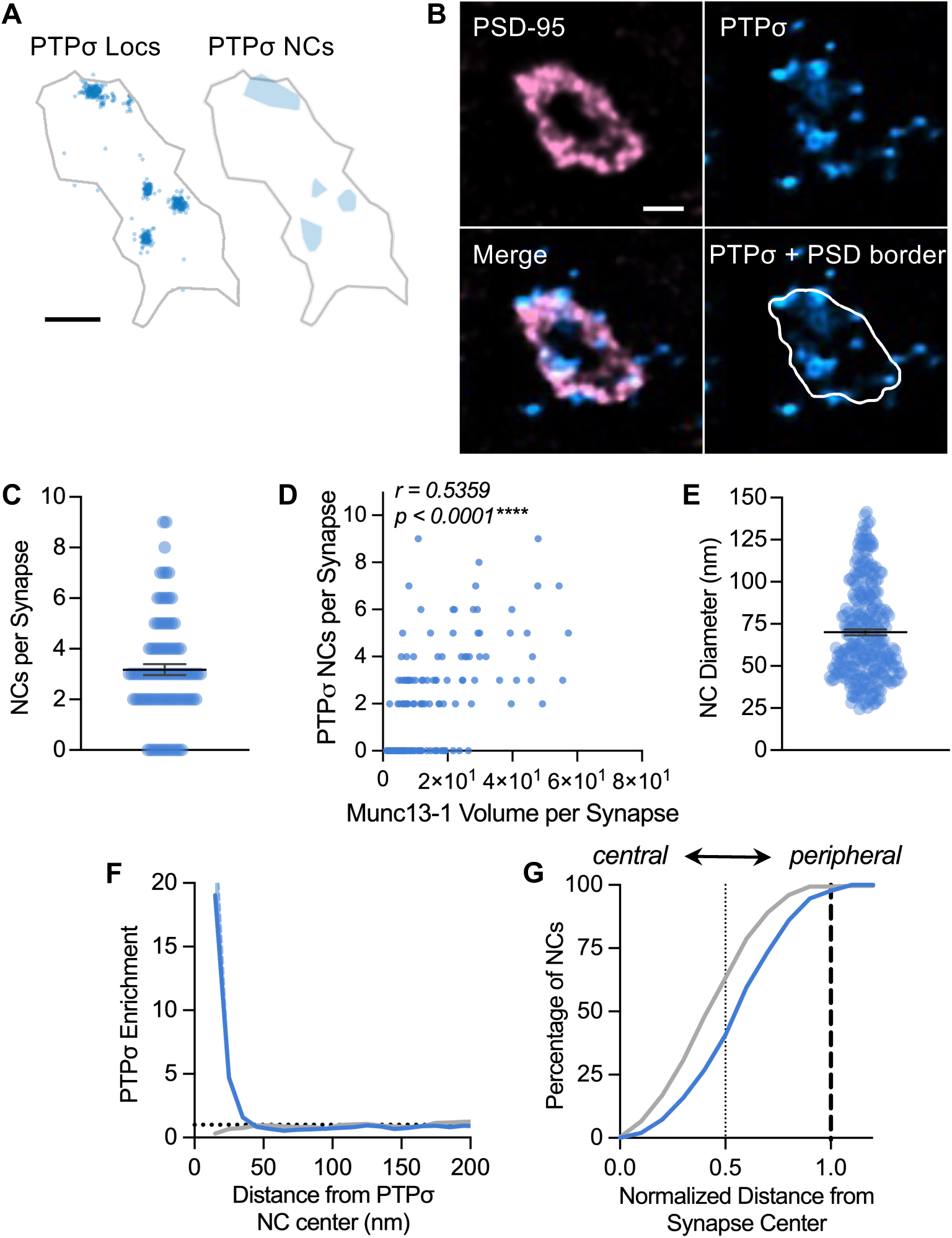
Endogenous PTPσ organizes into discrete NCs at excitatory synapses. **A)** Exemplar synapse showing DNA-PAINT localizations of PTPσ (blue) and the HDBSCAN detected NCs (blue shaded regions). Gray outline indicates PSD border, defined by PSD-95 localizations (not shown). Scale bar 130 nm. **B)** Exemplar 2D STED image of an excitatory synapse stained for PTPσ (blue) and PSD-95 (pink). NCs of PTPσ were visually apparent, consistent with **A**. Scale bar 250 nm. **C)** PTPσ organized into an average of 3 NCs per synapse. **D)** NC number was positively correlated with the volume of Munc13-1 localizations, a proxy for AZ volume (Pearson r = 0.532, R^2^=0.28, P<0.0001). **E)** The average PTPσ NC diameter was 70 ± 1.7 nm, when approximating a circular NC. **F)** Auto-enrichment analysis estimated a NC radius of 35-45 nm (blue) and a lack of enrichment around random positions (gray). Gray line indicates the enrichment around random positions. **G)** Cumulative distribution shows PTPσ NCs (blue line) were distributed more peripherally than if organized randomly within the same space (gray line). Data in **C**, **D and G** are individual synapses, and data in **E-F** and individual NCs. Lines in **C** and **E** are means ± SEM. Data in (**F**) are means ± SEM shading. N = 270 PTPσ NCs from 95 synapses. ****P<0.0001.

### PTPσ nanoclusters are offset from sites of vesicle release

Previous results suggest PTPσ may organize vesicle fusion machinery in the active zone by linking the RIM1/Munc13-1 complex to the plasma membrane (Emperador-Melero et al., 2024). In this model, one would expect PTPσ to be positioned nearby sites of vesicle fusion, rather than randomly. To test this, we expanded our analysis of this dataset to evaluate the spatial relationship of PTPσ NCs with Munc13-1 NCs (Fig. 4A), which mark putative sites of vesicle release (Böhme et al., 2016; Reddy-Alla et al., 2017; Sakamoto et al., 2018; Midorikawa et al., 2024). Consistent with previous results (Sakamoto et al., 2018; Dharmasri et al., 2023; Anderson et al., 2025), we observed on average 6.5 ± 0.3 Munc13-1 NCs per synapse (Fig. 4B). Munc13-1 NCs were more numerous than those of PTPσ, both on average across synapses (6.5 ± 0.3 Munc13-1 vs 3.2 ± 0.2 PTPσ NCs per synapse; Fig. 4B) and within single synapses (Fig. 4B). They were also slightly larger, both by HDBSCAN diameter (Fig. 4C) and AE analysis (Fig. 4D) (77.2 ± 1.2 Munc13-1 vs 70.0 ± 1.7 nm PTPσ). The number of PTPσ NCs scaled weakly but positively with the number of Munc13-1 NCs per synapse (r=0.342, *R^2^*=0.117, *p*<0.0001*; Fig. 4E). PTPσ NCs were positioned more peripherally than Munc13-1 NCs, which was equally likely to occupy the central and peripheral regions (Fig. 4F), suggesting the two proteins occupy partially overlapping regions of the synapse.

**Figure 4.**
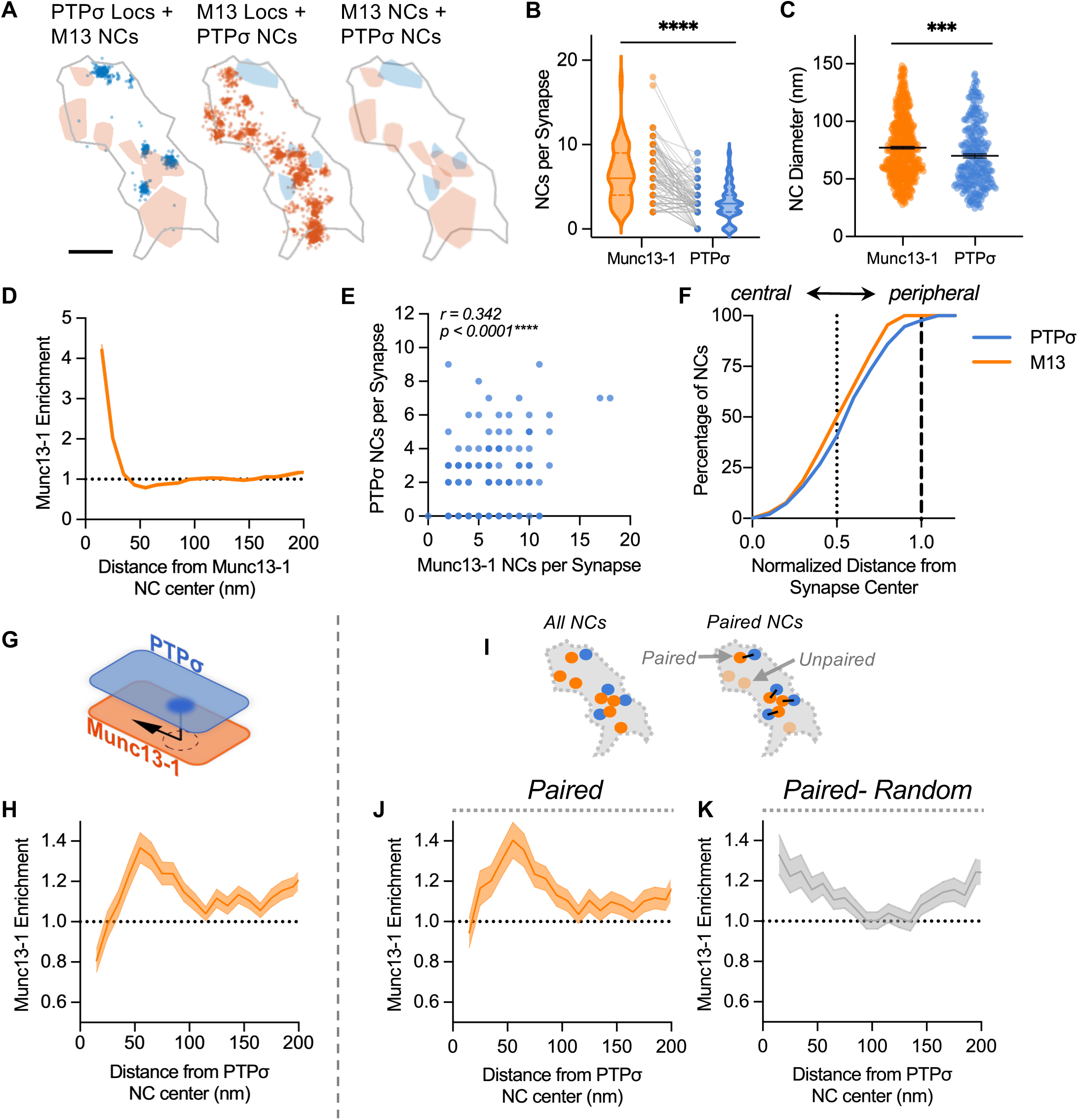
PTPσ NCs appear offset from Munc13-1 NCs. **A**) Exemplar synapse showing DNA-PAINT localizations of PTPσ (blue) and Munc13-1 (orange). Shaded regions indicate detected NCs of each protein overlaid with both the opposing protein’s localizations or detected NCs. Gray outline indicates PSD border, defined by PSD-95 localizations (not shown). Scale bar 130 nm. **B**) Munc13-1 NCs were more numerous per synapse (Wilcoxon matched-pairs signed rank test P<0.0001). Points (center) connected by grey line are individual synapses, with lines in violin (outside) at median ± IQR. **C**) Munc13-1 NCs were larger in diameter than PTPσ NCs (Mann-Whitney test P=0.0002). **D**) Munc13-1 auto-enrichment indicated a 35-45 nm NC radius, consistent with the estimated diameter in **C**. **E**) PTPσ NC number was weakly correlated with Munc13-1 NC number (Pearson r = 0.342, R^2^ = 0.117, P<0.0001). **F**) Cumulative distribution shows PTPσ NCs (blue) were distributed slightly more peripheral than Munc13-1 NCs (orange). **G**) Schematic representing the directionality of cross-enrichment analysis, radially measuring Munc13-1 protein enrichment (orange) from the center of PTPσ NCs (blue; NC dark blue). **H**) PTPσ NCs were, on average, de-enriched with Munc13-1 at distances <35 nm. **I**) Schematic of NC pairing to reveal stereotyped distances of closely associated NCs. Translucent circles in Paired condition indicate NCs lacking a mutually closest (“paired”) NC of the other protein. **J**) Pairing revealed the same de-enrichment of Munc13-1 within the first 35 nm of PTPσ NC center. **K**) Munc13-1 was enriched at the shortest distances from randomly positioned PTPσ NCs that were paired with a Munc13-1 NC by chance. Data in **D**, **H**, **J** and **K** are means ± SEM shading. Points in **C** are individual NCs, with lines at means ± SEM. n = 270 PTPσ and 566 Munc13-1 NCs from 95 synapses throughout, except (**J** and **K**) where n = 170 and 167 Munc13-1 NCs when paired with PTPσ NCs and random positions, respectively. ****P<0.0001.

To evaluate more specifically how Munc13-1 organizes with respect to PTPσ NCs, we performed a cross-enrichment analysis (Tang et al., 2016; Chen et al., 2020; Dharmasri et al., 2023), calculating the density of one protein as a function of distance from the other protein’s NC center (normalized to a randomized distribution of the protein being measured, Fig 4G). Munc13-1 was substantially de-enriched around the center of PTPσ NCs but became enriched at distances of 35 to 105 nm, with peak enrichment occurring between 50 and 60 nm away from a PTPσ NC center (Fig. 4H). This indicates that it is rare for Munc13-1 protein to be found within 35 nm of the center of a PTPσ NC. Notably, this distance is similar to the radius of a PTPσ NC; likewise, the distance range in which Munc13-1 becomes and stays enriched is quite comparable to the size of a Munc13-1 NC (∼70 nm vs ∼77 nm diameter, respectively). Therefore, PTPσ and Munc13-1 NCs appear to rarely occupy the same position in the synapse but instead lie adjacent to one another.

We examined the consistency of this offset to determine whether Munc13-1 is truly excluded from the center of PTPσ NCs. First, we repeated the cross-enrichment analysis on the subset of PTPσ and Munc13-1 NCs that were each other’s mutual nearest neighbors within a synapse (referred to as “paired NCs”; Fig. 4I). These NCs are the subset most closely positioned, so if the offset is systematic, then it should persist in this population. In this paired subset, NCs were still consistently offset from one another (Fig 4J), with the peak cross-enrichment at 50 to 55 nm, similar to the entire dataset (50 to 60 nm, Fig. 4H), indicating a lack of overlap. Second, we examined the cross-enrichment of Munc13-1 to random positions in the synapse when those positions were, by chance, mutually paired (Fig. 4I). This distribution was not offset from PTPσ NC center, peaking at the shortest distance (Fig. 4K), strongly suggesting that Munc13-1 is excluded from the vicinity of the peak of PTPσ NCs. Altogether, these data indicate the presence of a distinct offset between PTPσ and Munc13-1 NCs, consistent with their coordinated organization within the larger release site complex. This offset positioning between the NCs of two proteins in complex with one another could be explained by the required presence of intermediate proteins that mediate these interactions, for instance RIM1 or liprin-α3.

### PTPσ localizes to vesicle fusion sites regardless of their association to the nanocolumn

Given their mismatch in NC number, only a subset of Munc13-1 NCs have this systematic relationship with PTPσ, suggesting that these NCs might represent a unique class of release sites (Karlocai et al., 2021). Recent work has demonstrated that only a structurally distinct subset of Munc13-1 NCs are aligned trans-synaptically with PSD-95, i.e. are in the nanocolumn (Dharmasri et al., 2023; Anderson et al., 2025). Given its cis- and trans-synaptic interactors, PTPσ NCs could preferentially associate with this subset of Munc13-1 NCs. To identify nanocolumnar Munc13-1 NCs (“in nanocolumn” in Fig. 5), we expanded our analysis of this dataset to three proteins and identified Munc13-1 NCs that were significantly enriched with PSD-95 within 60 nm of the NC center (Tang et al., 2016; Anderson et al., 2025)(Fig. 5A), as well as those that were significantly de-enriched i.e. not in the nanocolumn. Using these NC assignments, we then paired PTPσ NCs or random positions with their mutually nearest Munc13-1 neighbor that was either in the nanocolumn or not, and calculated their distances and corresponding protein enrichments (Anderson et al., 2025). The distribution of distances between actual positions of PTPσ NCs and nanocolumnar Munc13-1 NCs was shifted rightward compared to distances from random positions (Fig. 5B), suggesting specific exclusion of PTPσ from nanocolumnar Munc13-1 NCs. Indeed, whether a Munc13-1 NC was associated with the nanocolumn or not made little difference to where mutually closest PTPσ NCs were positioned: the distances from PTPσ NCs was the same to both in and out of nanocolumn Munc13-1 NCs (Fig. 5C). When we subset cross-enrichments, regardless of whether a given Munc13-1 NC was in or out of the nanocolumn (Fig. 5E), there was an initial de-enrichment at position zero, which achieved enrichment at either 25 or 50 nm, respectively (Fig. 5E-F). These results support the conclusion that PTPσ NCs do not specifically organize near nanocolumnar release sites and instead either organize adjacent to release sites generally, or to release sites belonging to a different, as-yet-unknown structural or functional subset.

**Figure 5.**
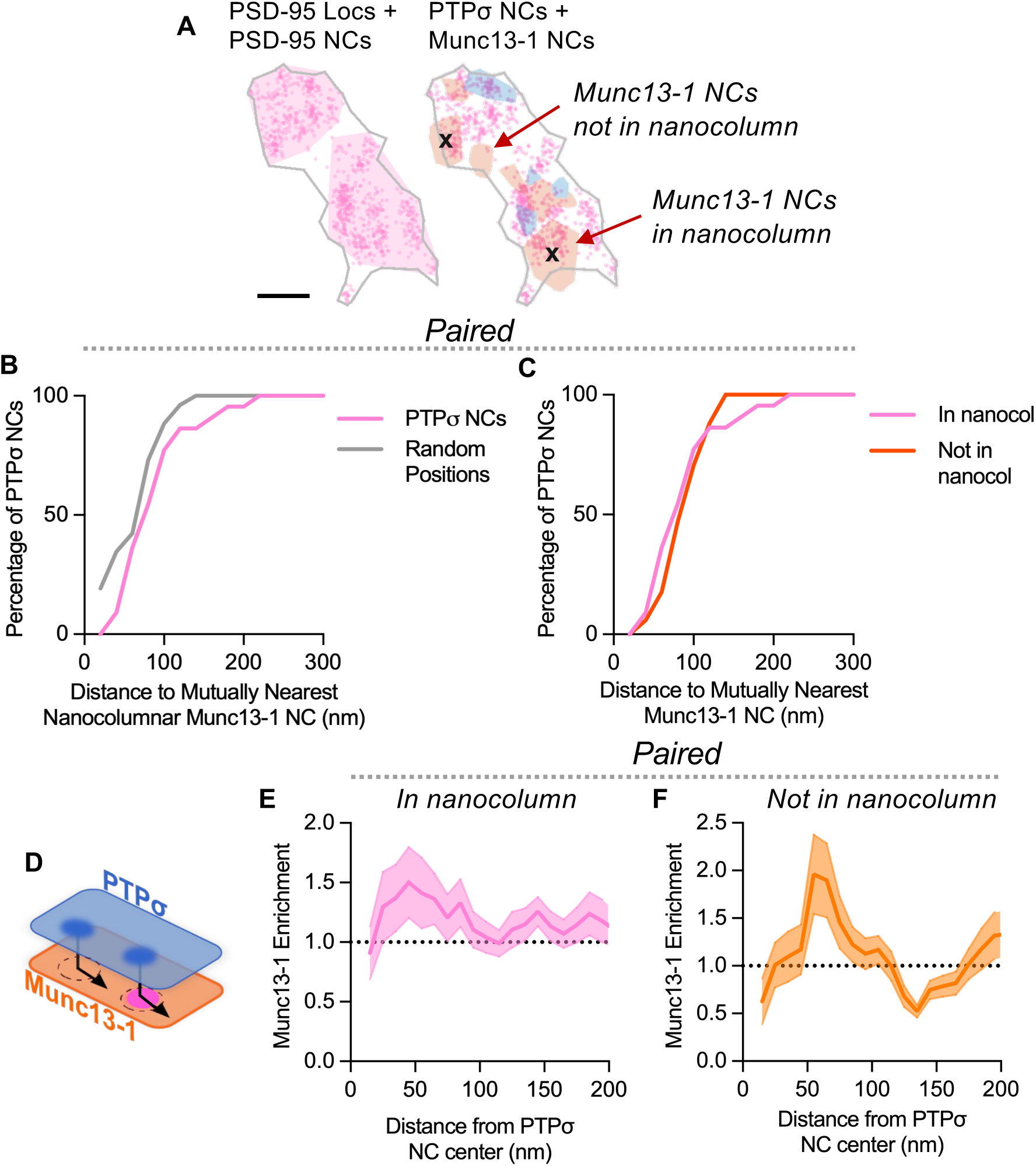
PTPσ localizes to vesicle fusion sites regardless of their association to the nanocolumn. **A**) Exemplar synapse showing DNA-PAINT localizations of PSD-95 (pink) and NCs (shading). Shaded regions representing PSD-95 (pink), PTPσ (blue), and Munc13-1 (orange) NCs are overlayed on PSD-95 localizations, with “x” marking locations of nanocolumnar Munc13-1 NCs. Gray outline indicates PSD border, defined by PSD-95 localizations. Scale bar 130 nm. **B**) Cumulative distribution shows PTPσ NCs are further from paired Munc13-1 NCs (as in **4I)** than if randomly positioned. **C**) Cumulative distribution shows PTPσ NCs are positioned at a similar distance away from a paired Munc13-1 NC regardless of whether that Munc13-1 NC is in the nanocolumn. **D**) Schematic demonstrating cross-enrichment analysis directionality, radially measuring Munc13-1 protein enrichment (orange) from the center of PTPσ NCs (blue; NC dark blue) that are paired with Munc13-1 NCs that are either in the nanocolumn (magenta) or not (pairing as in **4I**). **E**-**F**) The peak of PTPσ NC enrichment with Munc13-1 is similar regardless of whether the paired Munc13-1 NC was (**E**) in the nanocolumn or (**F**) not in the nanocolumn.

### NMDA receptors are offset from PTPσ NCs

Previous work suggests PTPσ could help regulate subsynaptic NMDAR positioning, as PTPσ KO reduces NMDAR currents without reducing NMDAR levels (Han et al., 2018; Sclip and Südhof, 2020b). PTPσ could play such a transsynaptic role through its cleft interactions with several proteins that bind NMDAR scaffolds (Dalva et al., 2000; Kwon et al., 2010; Takahashi and Craig, 2013; Ko et al., 2015; Li et al., 2015; Choi et al., 2016; Han et al., 2016; Farhy-Tselnicker et al., 2017), which could then influence NMDAR position. Therefore, one might expect PTPσ NCs to be systematically organized in respect to NMDARs. To test if this were the case, we evaluated the organization of GluN1 in respect to PTPσ NCs within the previously characterized set of synapses (Fig. 6A). GluN1 NCs were clearly apparent, consistent with previous reports (Kellermayer et al., 2018; Goncalves et al., 2020; Hruska et al., 2022). Compared to those of PTPσ, GluN1 NCs were more numerous (5.2 ± 0.4 GluN1 vs 3.2 ± 0.2 PTPσ NCs per synapse, respectively; Fig. 6B) and larger (88.1 ± 1.4 nm GluN1 vs 63.4 nm PTPσ; Fig. 6C-D). To evaluate their relationship to PTPσ, we first asked whether GluN1 and PTPσ NC numbers scaled together. We found no relationship between the number of PTPσ and GluN1 NCs per synapse (r= -0.076, *R^2^*= 0.006, *p*=0.464; Fig. 6E), unlike the relationship between PTPσ and Munc13-1 NCs which were weakly positively correlated (Fig. 4E). Despite this, PTPσ and GluN1 were similarly positioned closer to the edge of the synapse overall (Fig. 6F). This was especially apparent compared to Munc13-1, which was equally likely to occupy the central and peripheral regions (Fig. 6F). These data together suggest that GluN1 and PTPσ NCs occupy similar subregions of the synapse even though their abundance is not predictive of one another.

**Figure 6.**
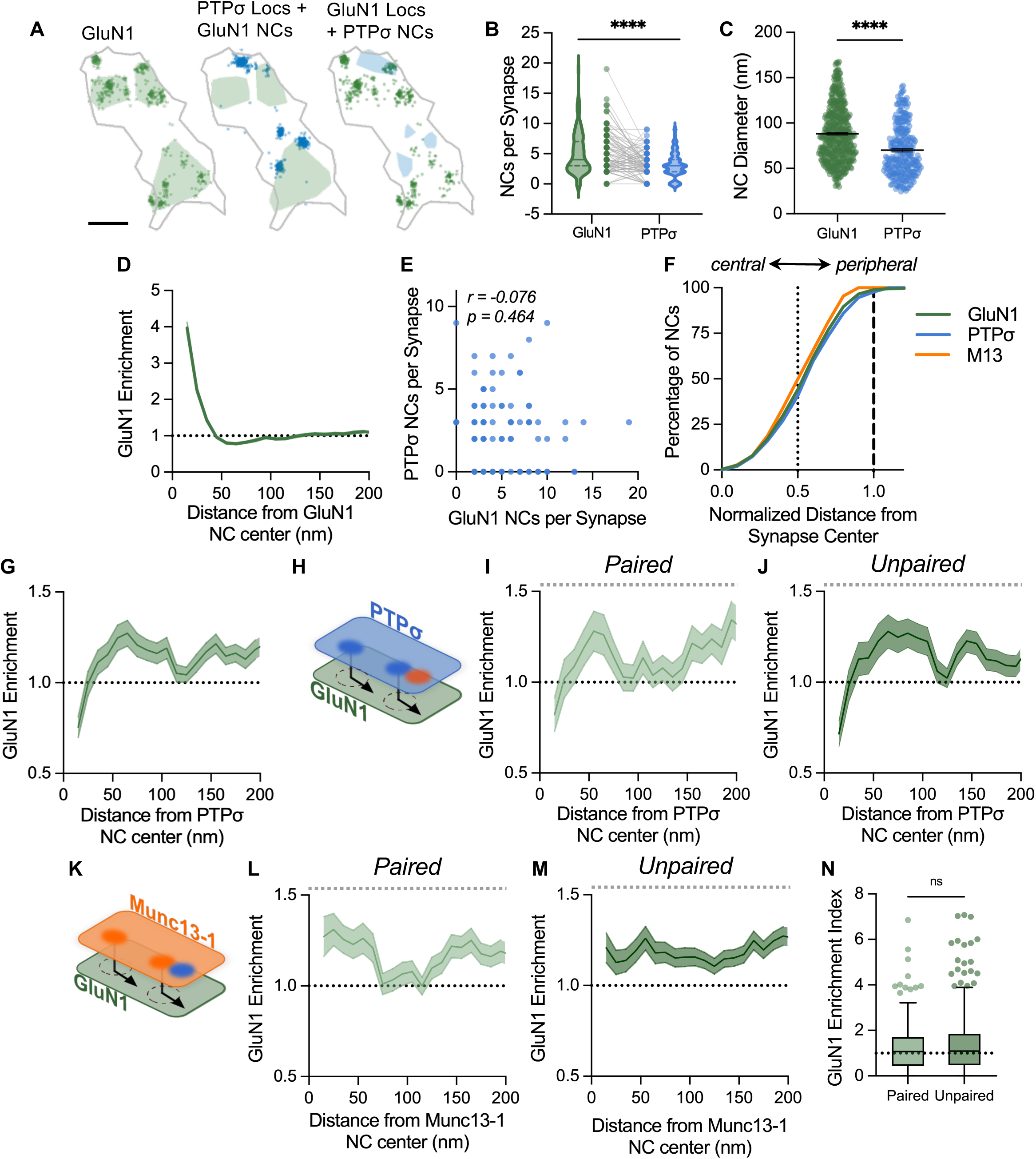
PTPσ NCs are systematically offset from GluN1 NCs. **A**) Exemplar synapse showing DNA-PAINT localizations of PTPσ (blue) and GluN1 (green). Shaded regions indicate detected NCs of each protein overlaid with both the opposing protein’s localizations or detected NCs. Gray outline indicates PSD border, defined by PSD-95 localizations (not shown). Scale bar 130 nm. **B**) GluN1 NCs were more numerous per synapse (Wilcoxon matched-pairs signed rank test P<0.0001). Points (center) connected by grey lines are individual synapses, with lines in violin (outside) at median ± IQR. **C**) GluN1 NCs were larger in diameter than PTPσ NCs (Mann-Whitney test P<0.0001). **D**) GluN1 auto-enrichment showed a 35-45 nm NC radius, consistent with the estimated diameter. **E**) PTPσ and GluN1 NC number per synapse were not correlated (Pearson r = -0.076, R^2^ = 0.005, P=0.464). **F**) Both GluN1 (green) and PTPσ (blue) NCs were distributed more peripheral than Munc13-1 (orange) NCs. **G**) Cross-enrichment analysis showed PTPσ NCs were, on average, de-enriched with GluN1 at distances <35 nm. **H**) Schematic demonstrating the cross-enrichment analysis directionality, radially measuring GluN1 protein enrichment (green) from the center of PTPσ NCs (blue; NC dark blue) that are or are not paired with their nearest neighboring Munc13-1 NCs (paired NC red; as in **4I**). **I-J**) PTPσ NCs that were paired (**I**) or unpaired (**J**) with Munc13-1 NCs were similarly de-enrichment with GluN1 within the first 35 nm (compare to **G**). **K**) Schematic demonstrating the cross-enrichment analysis directionality, radially measuring GluN1 protein enrichment (green) from the center of Munc13-1 NCs (orange; NC dark orange) that are or are not paired with their nearest neighboring PTPσ NCs (**dark blue;** same pairs as in **I-J**). **L**) When Munc13-1 NCs were paired with a PTPσ NC, GluN1 was enriched at short distances from the Munc13-1 NC center. **M**) Munc13-1 NCs that were not paired with a PTPσ NC had a consistently elevated enrichment of GluN1 at both short and far distances. **N**) There was no difference in GluN1 enrichment within the first 60 nm of Munc13-1 NCs with or without nearby PTPσ NCs. Data in **D**, **G**, **I**, **J**, **L** and **M** are means ± SEM shading. Points in **C** are individual NCs, with lines at means ± SEM. n = 270 PTPσ and 452 GluN1 NCs from 95 synapses throughout, except **I-J** where n = 100 and 169 PTPσ NCs when paired and unpaired with Munc13-1 NCs, respectively, and in **L-N**) where n = 179 and 312 Munc13-1 NCs when paired and unpaired with PTPσ NCs, respectively. ***P<0.001; ****P<0.0001.

We next measured the spatial relationship between GluN1 and PTPσ NCs using the cross-enrichment analysis. GluN1 density was initially de-enriched at the center of PTPσ NCs, but its enrichment increased and remained above random from 35 to 115 nm away from a PTPσ NC center (Fig. 6G). This relationship is strikingly similar to that observed for Munc13-1 density in relation to PTPσ NCs (Fig. 4H). However, it does not indicate whether the organization of these three proteins are dependent on one another. We tested the presence of this three-way enrichment in two ways. First, we separated the PTPσ NC enrichments with GluN1 by whether each PTPσ NC was mutually paired to a Munc13-1 NC or not (Fig. 6H). The stereotyped offset of GluN1 around PTPσ NCs was strikingly preserved regardless of whether PTPσ NCs had a neighboring Munc13-1 NC (Fig. 6I-J), indicating the spatial organization of PTPσ and GluN1 did not depend on the presence of a nearby SV fusion site. Second, because we know NMDARs localize to only a subset of SV fusion sites (Li et al., 2021; Hruska et al., 2022; Anderson et al., 2025), we next assessed whether GluN1 was differentially organized around the subset of Munc13-1 NCs that were paired or not with a PTPσ NC (Fig. 6K). Munc13-1 NCs with a nearby, paired PTPσ NC were enriched with GluN1 from near their center to ∼70 nm away, before enrichment decreased to random (Fig. 6L). By contrast, Munc13-1 NCs *without* a nearby PTPσ NC had a different GluN1 enrichment profile, where GluN1 enrichment was essentially flat across distance (Fig. 6M). The complexity of these curves is likely representative of the great range in distances over which GluN1 can be found opposite Munc13-1 NCs. One interpretation of the differences between these curves could be that PTPσ, through its transsynaptic interactions, plays a role in refining the position of GluN1 around SV fusion sites. However, we found that PTPσ did not predict GluN1 enrichment to SV fusion sites in general, as the average enrichment of GluN1 near (within 60 nm) a Munc13-1 NC center was not significantly different depending on whether a PTPσ NC was nearby (Fig. 6N). Together, these analyses suggest that these three proteins organize in a stereotyped manner but largely independent from their nanocolumn relationship.

While PTPσ itself was not selectively localized to the nanocolumn, it is still possible that its presence at nanocolumn sites will influence GluN1 organization opposite to it. To evaluate this possibility, we integrated all four proteins from our synapse dataset and performed a cross-enrichment analysis to assess the density of GluN1 opposing PTPσ NCs that were or were not paired with nanocolumnar Munc13-1 NCs (Fig. 7A). Both relationships (Fig. 7B-C) recapitulated the previously described offset (Fig. 6G) of GluN1 density from PTPσ NC centers. The curves were largely comparable, though there was a small apparent shift of the enrichment’s peak from ∼30 nm away when unpaired to ∼50 nm away when PTPσ NCs were paired with the nanocolumn (Fig. 7B-C). Reciprocally, we asked whether the enrichment of NMDARs at the nanocolumn is influenced by the presence of nearby PTPσ NCs. The resulting cross-enrichment curves analyzing the density of GluN1 opposite nanocolumn-associated Munc13-1 NCs that were paired or not with a nearby PTPσ NC (Fig. 7D) were noisier due to the increasingly stringent requirements and sub-setting necessary. Nevertheless, GluN1 enrichment was similar regardless of the presence of PTPσ NCs, showing a weak initial enrichment of GluN1 opposing the center of nanocolumnar Munc13-1 NCs decaying similarly with distance from the center of the nanocolumn (Fig. 7E-F). Additionally, the average enrichment of GluN1 within the first 60 nm of a given nanocolumnar Munc13-1 NC center was not different when paired with a neighboring PTPσ NC (Fig. 7G). While these data support previous work demonstrating the nanocolumn localization of NMDA receptors (Anderson et al., 2025), they also demonstrate this positioning likely occurs regardless of their association with PTPσ NCs.

**Figure 7.**
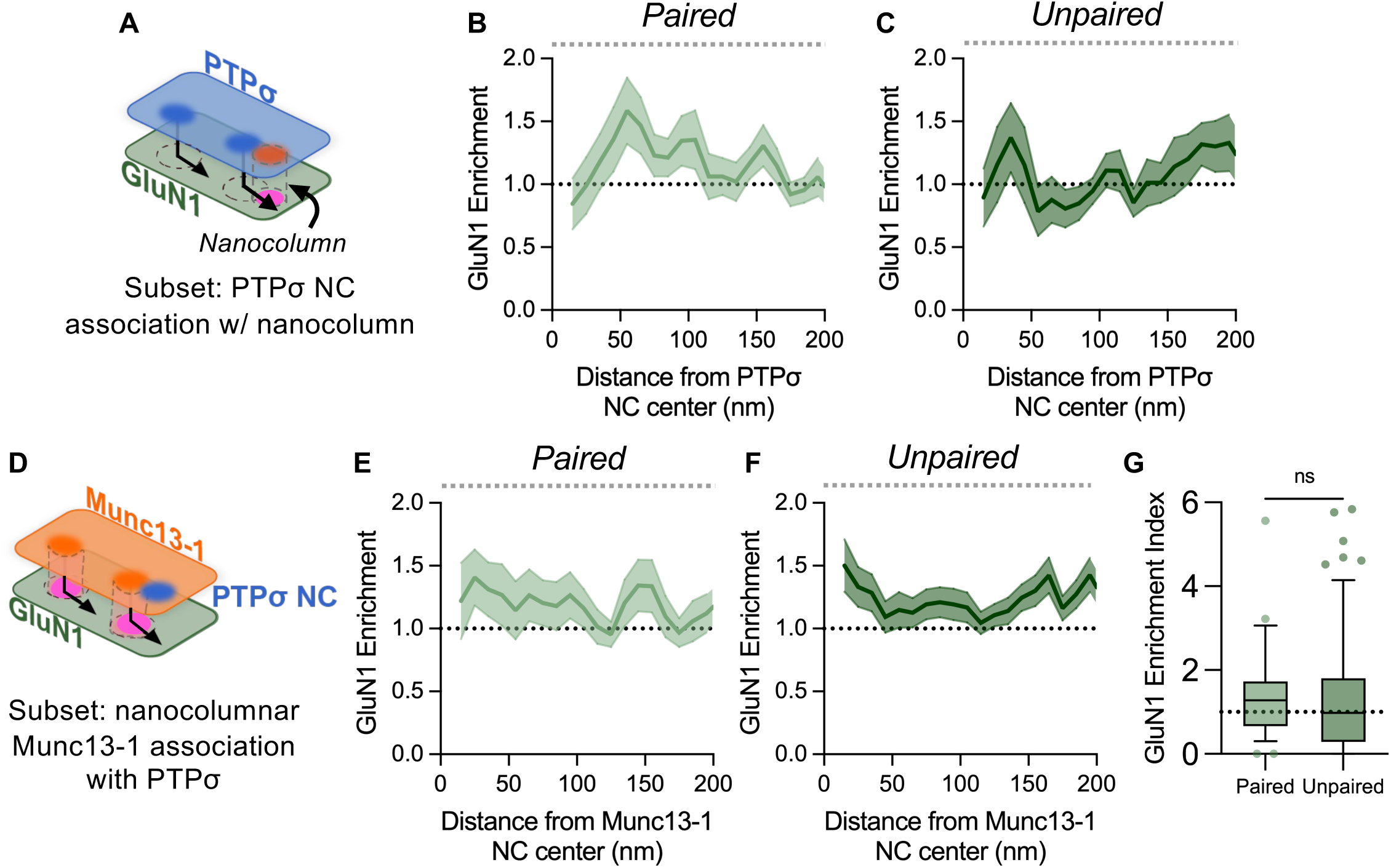
The presence of PTPσ does not increase GluN1 enrichment at the nanocolumn. **A**) Schematic demonstrating cross-enrichment analysis directionality, radially measuring GluN1 protein enrichment (green) from the center of PTPσ NCs (blue; NC dark blue) that are paired or unpaired with their nearest neighboring nanocolumnar Munc13-1 NCs (paired Munc13-1 NC are orange, aligned PSD-95 NCs are magenta; pairing from **4I**). **B**-**C**) PTPσ NCs that were (**B**) paired or (**C**) not paired with nanocolumnar Munc13-1 NCs were similarly enriched with GluN1, with a shift in the peak of GluN1 enrichment. **D**) Schematic demonstrating cross-enrichment analysis directionality, radially measuring GluN1 protein (green) enrichment from the center of nanocolumnar Munc13-1 NCs (orange; NC dark orange; aligned PSD-95 NCs are magenta) that are either paired or unpaired with their nearest neighboring PTPσ NCs (dark blue). **E**-**F**) Nanocolumnar Munc13-1 enrichment with GluN1 was not significantly different regardless of whether the Munc13-1 NC was (**E**) paired or (**F**) unpaired with a neighboring PTPσ NC.

Altogether, the intricate spatial relationships among these four proteins reveal that presynaptic PTPσ appears likely to have only minor influence on NMDAR positioning with respect to glutamate release sites while still occupying a distinct position around release sites themselves.

## Discussion

We used confocal, STED and single-molecule localization microscopy to investigate endogenous PTPσ expression and subcellular localization to evaluate a potential role for this important molecule in coordinating synapse nanostructure. We found PTPσ was preferentially expressed in the presynaptic compartment of established excitatory synapses and further evaluated its unique nanoscale organization within these structures. Our data reveal novel subsynaptic relationships with Munc13-1 within the AZ and with NMDARs trans-synaptically, suggesting PTPσ may play one or more unexpected roles.

### Munc13-1 offset and release site architecture

A central aspect of our data provides new insights to the organization of the AZ and the SV fusion sites within it. PTPσ has previously been implicated in SV fusion as its ablation reduces release probability (Han et al., 2018, 2020; Sclip and Südhof, 2020a). Furthermore, PTPσ directly binds liprin-α3 (Wakita et al., 2020) and is indirectly linked to RIM1 through shared interactions with SV priming machinery (Wong et al., 2018; Emperador-Melero et al., 2024; Jin et al., 2025). Together, these functional and structural relationships led to our prediction that PTPσ would localize to sites of vesicle fusion. However, though the position of PTPσ in the AZ was indeed related to SV fusion sites marked by Munc13-1 NCs (Böhme et al., 2016; Reddy-Alla et al., 2017; Sakamoto et al., 2018), we found surprisingly that NCs of PTPσ were consistently offset from, or adjacent to, Munc13-1 NCs. This juxtaposed organization suggests PTPσ may participate in vesicle fusion-related processes through close spatial coupling to release sites despite not residing within the fusion machinery complex itself. Recent cryo-ET studies provide possible structural rationale for this, with visualization of cleft-spanning filaments linking presynaptic vesicles with PSD material (Martinez-Sanchez et al., 2021; Cole and Reese, 2023) as well as membrane-proximal protein densities in the AZ displaced ∼50 nm from the nearest docked synaptic vesicle (Held et al., 2024). Though the protein composition of these nearby densities has not been determined, our data suggest PTPσ may be a component.

Several lines of evidence suggest a new model of protein organization around SV fusion sites that enables these proteins to be spatially segregated while also acting in complex (Fig. 8). First, we note recent evidence that Munc13-1 forms membrane-associated hexameric rings that aid vesicle priming and that SNARE-mediated fusion occurs in the center of this ring (Quade et al., 2019; Grushin et al., 2022; Li et al., 2023; Eilts et al., 2026). While the specific nature and generalized presence of such a ring structure remain under investigation, we consider it likely that Munc13-1 NCs visualized here represent multi-protein structure surrounding a fusion site. In this case, crowding cause by the presence of the docked SV and SNAREs may limit close access of many other proteins. Consistent with this, RIM1 (Eilts et al., 2026) was consistently positioned outside of hexameric, ring-shaped Munc13-1 structures (presumably NCs) visualized in native synapses using a combination of tissue expansion and dSTORM. Additionally, the multitude of cis- and trans-synaptic interactions made by PTPσ may provide further steric hindrance, limiting its close approach to SV fusion sites (Held et al., 2024). Thus it may be a combination of both the indirect nature of PTPσ’s interaction with RIM1 and steric hindrance at the plasma membrane that positions PTPσ at this stereotyped distance away from SV fusion sites and the surrounding protein machinery (Fig. 8) (Wakita et al., 2020; Emperador-Melero et al., 2024; Jin et al., 2025). Considering PTPσ directly binds liprin-α3 (Wakita et al., 2020) and forms protein condensates with liprin-α3 and RIM1 in heterologous cells (Emperador-Melero et al., 2024), it is possible that these interactions could further support its peripheral localization. It will be important in the future to evaluate the subsynaptic organization of liprins in respect to these proteins as well as their liquid-liquid phase-separation status in the AZ.

**Figure 8.**
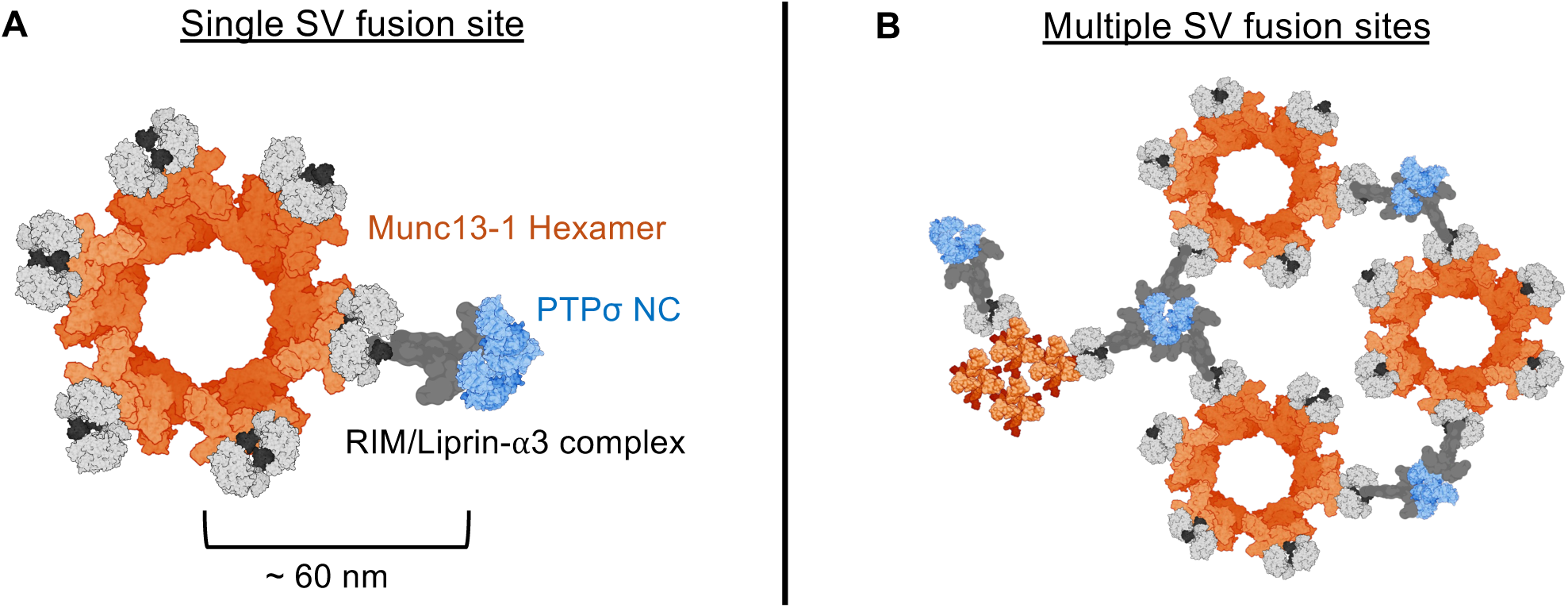
Prospective working model of how PTPσ may be positioned ∼60 nm from Munc13-1 NCs while still functioning in complex with vesicle fusion machinery.

Emerging from this model is the general notion that even AZ proteins positioned seemingly distant from SV fusion sites may still function in concert to support SV fusion. In the case of PTPσ, these functions may be mediated persistently via intervening protein interactions (i.e. PTPσ-liprin-RIM-Rab3a/SV), or could be dynamic, perhaps by aiding Munc13-1 in its initial membrane tethering interactions before Munc13-1 laterally engages with other machinery to proceed in the priming process (Quade et al., 2019; Held et al., 2024). Additional advances to combine high-resolution techniques for specific protein labeling such as DNA-PAINT with ultrastructural identification (e.g. cryo-ET) will be required to better understand how the spatial positions of proteins within this complex fundamentally support vesicle fusion.

### PTPσ association with a subset of fusion sites

Beyond the positional offset from release sites, we also observed more Munc13-1 NCs per synapse than PTPσ, suggesting PTPσ could preferentially organize with a specific subset of fusion sites. Indeed, there is significant heterogeneity in both the function and protein composition of SV fusion sites within the same synapse (Akbergenova et al., 2018; Maschi and Klyachko, 2020; Karlocai et al., 2021; Aldahabi et al., 2022; Gou et al., 2022; Emperador-Melero et al., 2024; Anderson et al., 2025), and functional data suggest evoked and spontaneous fusion occur at different subsets of release sites (Kavalali, 2015; Yang et al., 2025). While we predicted a role for PTPσ NCs in evoked SV fusion, we did not find a differential organization of PTPσ around SV fusion sites in the nanocolumn, where evoked fusion preferentially occurs (Tang et al., 2016; Biederer et al., 2017; Haas et al., 2018; Ramsey et al., 2021; Muttathukunnel et al., 2022) (but see also Held 2024). Thus, it is possible that PTPσ could unexpectedly regulate spontaneous transmission. In support of this, PTPσ NCs tended to localize toward the synapse edge, consistent with the observed positioning of spontaneous fusion events (Kavalali, 2015; Tang et al., 2016). However, the structural features corresponding to spontaneous fusion sites remain unknown, limiting our ability to isolate that population from our current dataset. Altogether, PTPσ’s organization at the AZ to a subset of fusion sites is consistent with a model where the AZ is composed of multiple interdigitated nanodomains that dictate unique functional sub-regions of the synapse (Guzikowski and Kavalali, 2021; Metzbower et al., 2024).

### Subsynaptic PTPσ organization and phosphatase function

Because PTPσ is a phosphatase, nanoclustering of the protein may have functional impact beyond the structural roles mediated by its protein interactions. PTPσ expression alters the phosphorylation status of many presynaptic proteins, including CASKIN2, vGluT2, synaptotagmin-11, CACNA1A, APP and Bassoon (Kim et al., 2020; Han et al., 2025), though interestingly there is not evidence of it regulating critical members of SV fusion machinery such as RIM1 or Munc13. Intriguingly, when LAR-RPTPs cluster on the cell surface their phosphatase activity is inhibited (Bilwes et al., 1996; Majeti et al., 1998; Meng et al., 2000; Xie et al., 2020). This suggests PTPσ NCs may create biochemically distinct subdomains of the AZ representing weakened points of phosphatase activity around particular SV fusion sites. It is possible that in the absence of its phosphatase activity, PTPσ could serve a more structural role in bridging transcellular compartments. In concert with the offset, these data suggest a broader architecture for functional regions within the synapse.

### NMDAR offset and positioning at fusion sites

PTPσ ablation reduces evoked NMDAR amplitudes (Kim et al., 2020; Sclip and Südhof, 2020a; Han et al., 2025), suggesting PTPσ may play a trans-synaptic role in NMDAR regulation. We therefore hypothesized that PTPσ may function to subsynaptically position NMDARs within specific regions of the postsynaptic density. In this model, PTPσ NCs would be aligned with NMDAR NCs to physically position them across the synapse. Though we found that NCs of PTPσ and GluN1 were similarly distributed over the radial extent of the synapse, their NCs were not trans-synaptically aligned. Instead, PTPσ NCs appeared substantially offset from GluN1 NCs, similar to their relationship to SV fusion sites. Because PTPσ is not known to bind NMDARs directly, such an offset is not surprising. Indeed, there are a variety of molecules that could mediate indirect interactions between PTPσ and NMDARs and establish this offset relationship, including postsynaptic cell-adhesion molecules (Dalva et al., 2000; Kwon et al., 2010; Budreck et al., 2013; Li et al., 2015; Choi et al., 2016; Han et al., 2016) or additional postsynaptic or cleft-resident proteins (Ko et al., 2015; Farhy-Tselnicker et al., 2017; Liakath-Ali et al., 2022).

Altogether, these data illustrate the power of multiplexed nanoimaging to reveal the intricate spatial relationships of proteins within synapses. We conclude that the positioning of PTPσ is related not only to active zone proteins as expected based on its presynaptic occupancy and known roles, but also to NMDARs. These relationships raise the possibility that PTPσ may play multiple roles to fine-tune the functional performance of synapses. Presynaptically, its occupancy at a subset of release sites suggests it may shape the spatial distribution of primed SVs across the face of the synapse. Postsynaptically, it may fine tune NMDAR positioning through intermediate interactors. Particularly in concert, such effects are likely to modulate NMDAR synaptic signaling with implications for developmental and activity-dependent modulation of neural circuit function. Based on discerning the pattern of PTPσ positioning, this work thus will help direct the design of future experiments to test specific functional roles.

## Acknowledgements

This work was supported by an MPower Early Scholars Investment Fund award to EMD and the National Institutes of Health (F31-NS129086 to EMD, T32-GM159581 to SM, R37-MH080046 to TAB, R21-MH141559 to TAB & ADL). STED images were acquired in the University of Maryland School of Medicine Center for Innovative Biomedical Resources Core Confocal Facility, with support from the University of Maryland Medicine Institute for Neuroscience Discovery (UM-MIND). We thank Minerva Contreras for outstanding technical support, members of the Blanpied lab for important discussions and evaluation of the manuscript, and members of the Jungmann lab for their assistance with DNA-PAINT optimization.

---

**Supplemental Figure 1.**
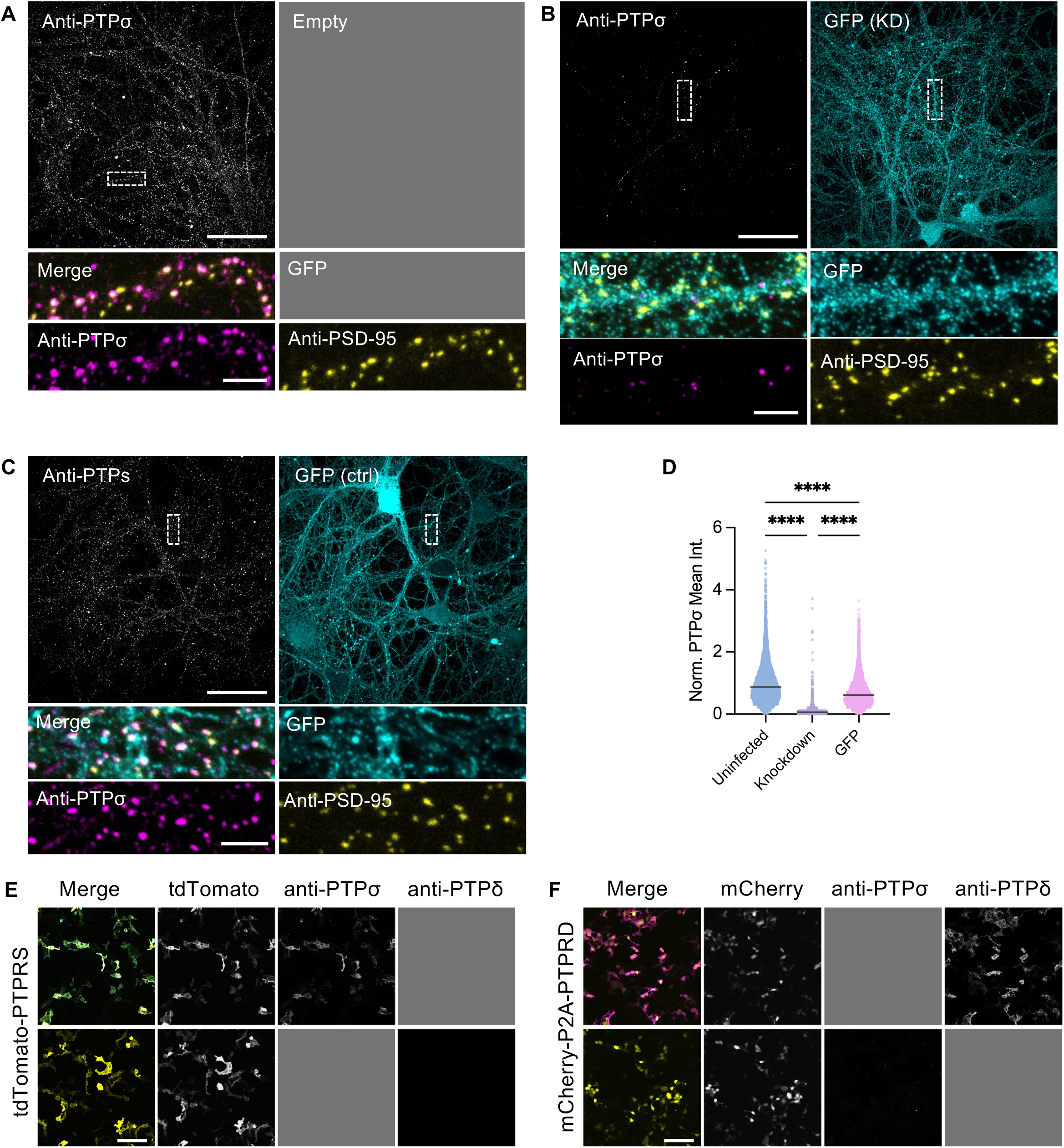
Confirmation of anti-PTPσ specificity in DIV21 rat primary hippocampal neuron culture. Immunolabeling of PTPσ (magenta) and PSD-95 (yellow) with or without a GFP cell fill (cyan, viral reporter) in (**A**) uninfected, (**B**) PTPσ knockdown-infected or (**C**) control GFP cell fill-infected DIV21 rat primary hippocampal neurons. Dashed boxes indicate location of insets to right of overviews. Overview shows synaptic staining on one side and GFP cell fill on other. Scale bar 50 and 5 µm inset. **D**) Normalized PTPσ mean intensity within PSD-95 ROIs is significantly reduced when PTPσ is knocked down, compared to uninfected or control GFP cell fill-infected controls. Points in **D** are individual PSD-95 ROIs. Specificity validation of anti-PTPσ. **E**) Anti-PTPσ labeled PTPσ, but not PTPδ, protein overexpressed in HEK cells. Scale bar 50 µm. **F**) Anti-PTPσ does not label PTPδ protein overexpressed in HEK cells, while anti-PTPδ does. Scale bar 50 µm. Gray boxes are present to indicate channels not applicable for that immunostaining.

